# Interferon-Induced Transmembrane Proteins Inhibit Infection by the Kaposi’s Sarcoma-Associated Herpesvirus and the Related Rhesus Monkey Rhadinovirus in a Cell Type-Specific Manner

**DOI:** 10.1101/2021.07.16.452764

**Authors:** Bojan F. Hörnich, Anna K. Großkopf, Candice J. Dcosta, Sarah Schlagowski, Alexander S. Hahn

## Abstract

The interferon-induced transmembrane proteins (IFITMs) are broad-spectrum antiviral proteins that inhibit the entry of enveloped viruses. We analyzed the effect of IFITMs on the gamma2-herpesviruses Kaposi’s sarcoma-associated herpesvirus (KSHV) and the closely related rhesus monkey rhadinovirus (RRV). We used CRISPR/Cas9-mediated gene knockout to generate A549, human foreskin fibroblast (HFF) and human umbilical vein endothelial cells (HUVEC) with combined IFITM1/2/3 knockout and identified IFITMs as cell type-dependent inhibitors of KSHV and RRV infection in A549 and HFF but not HUVEC. IFITM overexpression revealed IFITM1 as the relevant IFITM that inhibits KSHV and RRV infection. Fluorescent KSHV particles did not pronouncedly colocalize with IFITM-positive compartments. However, we found that KSHV and RRV glycoprotein-mediated cell-cell fusion is enhanced upon IFITM1/2/3 knockout. Taken together, we identified IFITM1 as a cell type-dependent restriction factor of KSHV and RRV that acts at the level of membrane fusion. Strikingly, we observed that the endotheliotropic KSHV circumvents IFITM-mediated restriction in HUVEC despite high IFITM expression, while influenza A virus (IAV) glycoprotein-driven entry into HUVEC is potently restricted by IFITMs even in the absence of interferon.

**IMPORTANCE:** IFITM proteins are the first line of defense against infection by many pathogens, which may also have therapeutic importance, as they, among other effectors, mediate the antiviral effect of interferons. Neither their function against herpesviruses nor their mechanism of action are well understood. We report here that in some cells, but not in, for example, primary umbilical vein endothelial cells, IFITM1 restricts KSHV and RRV, and that, mechanistically, this is likely effected by reducing the fusogenicity of the cell membrane. Further, we demonstrate potent inhibition of IAV glycoprotein-driven infection of cells of extrapulmonary origin by high constitutive IFITM expression.

## INTRODUCTION

The family of interferon-induced transmembrane proteins (IFITMs) are small membrane proteins that exhibit antiviral activity toward a broad variety of viruses (1–6). There are five IFITMs present in the human genome, but only IFITM1, IFITM2, and IFITM3 are known to be immune-related and interferon(IFN)-inducible (reviewed in (6) and (7)). IFITM1 localizes to the plasma membrane, while IFITM2 and IFITM3 localize to endosomes/lysosomes (5, 8). The exact mechanism of IFITM-mediated restriction of viral replication is not completely understood. It is, however, clear that restriction mainly occurs at the viral entry stage (3, 9, 10). According to some reports, IFITMs modify the overall membrane fusogenicity, by modification of the membrane lipid composition and/or the membrane rigidity and thus prevent virus-host membrane fusion (11–14), probably causing arrest of the fusion pore opening following hemifusion (11, 12, 15). Other modes of action such as e.g. recruitment of additional antiviral factors, altered endocytic trafficking or interference with vacuolar ATPase have been postulated as well (reviewed in (16)).

The majority of IFITM-restricted viruses are RNA viruses. The interplay of IFITMs with DNA-viruses has been studied less extensively and with more ambiguous results. While vaccinia virus and herpes simplex virus (HSV-1) are restricted by overexpression of individual IFITM proteins (17, 18), human papillomavirus 16 (HPV16) and the non-enveloped adenovirus type 5 are not (19). Interestingly, for the human cytomegalovirus (HCMV), small interfering RNA (siRNA)-mediated IFITM knockdown resulted in reduced infection and disturbed virus assembly (20). Varying results were obtained for Epstein-Barr virus (EBV), a gammaherpesvirus. While the initial entry of EBV was enhanced by overexpression of IFITM1 (21, 22), incorporation of IFITM2/3 into viral particles reduced the infectivity of progeny virus, whereas IFITM1 incorporation had no effect (23). Together, the literature on IFITM-mediated effects on the alphaherpesvirus HSV-1, the betaherpesvirus HCMV, and the gammaherpesvirus EBV indicate differences in the activity of IFITM proteins toward different herpesvirus subfamilies. The Kaposi’s sarcoma-associated herpesvirus (KSHV) and the related rhesus monkey rhadinovirus (RRV) belong to the *γ*-herpesvirus subfamily (24). KSHV is associated with Kaposi’s sarcoma (KS), multicentric Castleman’s disease, primary effusion lymphoma (reviewed in (25)), osteosarcoma (26), and KSHV inflammatory cytokine syndrome (KICS) (27). The incidence of KSHV-related disease and KSHV seroprevalence are low in industrial countries (28, 29), but KSHV represents a significant health burden in Sub-Saharan Africa, where KSHV-related cancers are common (30, 31).

KSHV and RRV exhibit broad cell tropism *in vitro* (32, 33). Both viruses encode for a set of glycoproteins (g) that mediate entry and are conserved among herpesviruses. Of these gH, gL, and gB are the most extensively studied (reviewed in (34)). KSHV and RRV enter many cell types through the interaction of the gH/gL complex with members of the Ephrin receptor tyrosine kinase family (Ephs) (35–37) and, in case of RRV also with members of the Plexin domain-containing protein family (38). KSHV also interacts with heparan sulfate and integrins (39–41). Entry of both viruses mainly occurs via endocytotic routes (33, 42–44). Following internalization, the viral membrane fuses with the host membrane. Several reports implicate the gH/gL complex together with gB as the minimal set of glycoproteins required for membrane fusion (45, 46, 38). One study reported an enhancing role of IFITMs in the infection of the BJAB B cell line and human dermal microvascular endothelial cells (HMVEC-D) cells by KSHV, EBV, and herpes simplex virus 2 (HSV-2) (21). However, given the considerable differences between KSHV and RRV entry into B cells and different adherent cells (33, 36, 37), in particular, as KSHV infection of B cell lines is, with a few exceptions, only efficient through cell-to-cell transfer (47–49), we hypothesized that IFITM-mediated restriction may be dependent on the type of target cell. Another question that we sought to address is whether IFITMs restrict RRV in human cells.

## RESULTS

### KSHV induces IFITM expression in A549 cells

We first validated the specificity of the antibodies used in this study for Western blot analysis after directed expression (Fig. 1 A). Next, we examined expression of IFITM proteins at baseline levels and after stimuli such as virus infection in the human lung epithelial cell line A549, which has been well characterized with regard to IFN signaling and IFITM expression (50–53). We infected the cells with KSHV BAC16 recombinant virus carrying a green fluorescent protein (GFP) reporter gene, and RRV-YFP carrying a yellow fluorescent protein (YFP) reporter gene (Fig. 1 B). Treatment with H_2_O and IFN-*α* served as negative and positive controls for IFITM induction, respectively. IFITM2 and IFITM3 were detected at low levels without IFN treatment, while IFITM1 and human myxovirus resistance gene 1 (MxA), another IFN-induced protein, were not detectable without stimulation (Fig. 1 C). At the 1 h timepoint, neither treatment induced IFITM or MxA expression relative to the background. At the 24 h timepoint, induction over background levels of IFITM1, IFITM2, IFITM3, and MxA was observed in IFN-*α*-treated or KSHV-infected cells, but not in RRV-infected cells. At 48 h, IFITM3 was also slightly induced by RRV, and IFITM2 induction relative to H_2_O treatment was barely discernable anymore. Basal IFITM expression also increased slightly over time after plating. In summary, KSHV-containing inoculum and IFN-*α* induced IFITM expression.

**Figure 1.**
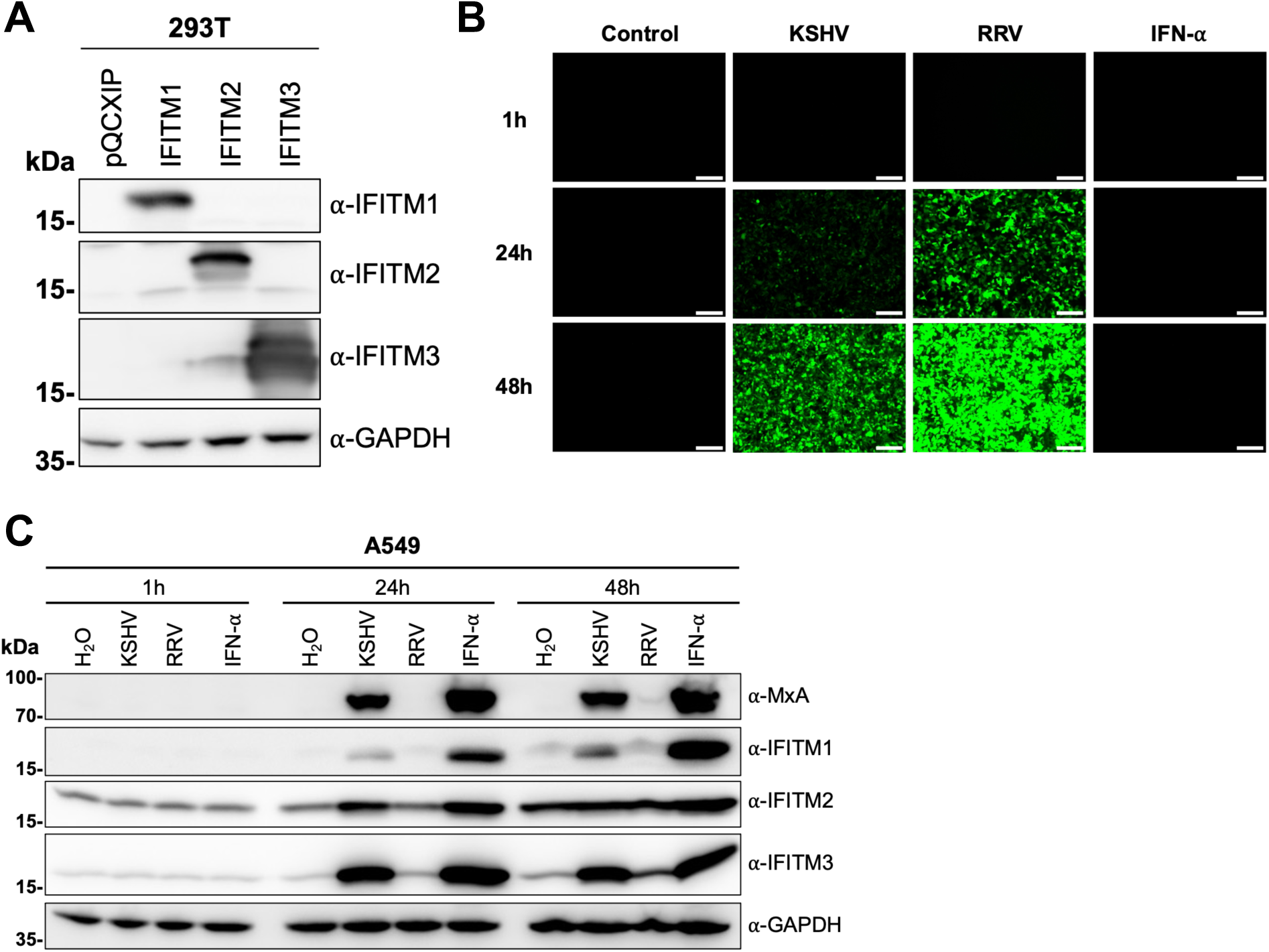
KSHV induces IFITM1, IFITM2 and IFITM3 expression in A549 cells. **(A)** Western blot of 293T cells transduced with pQCXIP-constructs to express IFITM1-3 or pQCXIP (empty vector). IFITMs were detected using the respective IFITM-antibody, GAPDH served as loading control. **(B)** Fluorescence microscopy images (scale 200µm) and **(C)** Western blot analysis of A549 cells infected with KSHV-GFP or RRV-YFP or treated with H_2_O or IFN-α (5000 U/ml) for the indicated time and harvested using SDS sample buffer. IFITM expression was detected with antibodies shown in (A). MxA served as control for IFN-stimulated gene induction; GAPDH served as loading control.

### Triple knockout of IFITM1/2/3 enhances KSHV and RRV infection of A549 and HFF

Overexpression of IFITMs alters their subcellular localization ((6), own observations), IFITMs are usually induced together, and recent studies report that IFITMs form homo- and hetero-oligomers (54–56) and might thus act synergistically. We therefore used CRISPR/Cas9 to generate triple IFITM1/2/3 knockout cells to study the effects of basal IFITM expression as well as IFN-induced IFITM expression on KSHV and RRV infection. We identified two single guide RNAs (sgRNAs) (sgIFITM1/2/3-a, sgIFITM1/2/3-b), which target the second exon of all three immune-related IFITMs (Fig. 2 A, 2 B). These sgRNAs were transduced together with Cas9 using the lentiCRISPRv2 system (57).

**Figure 2.**
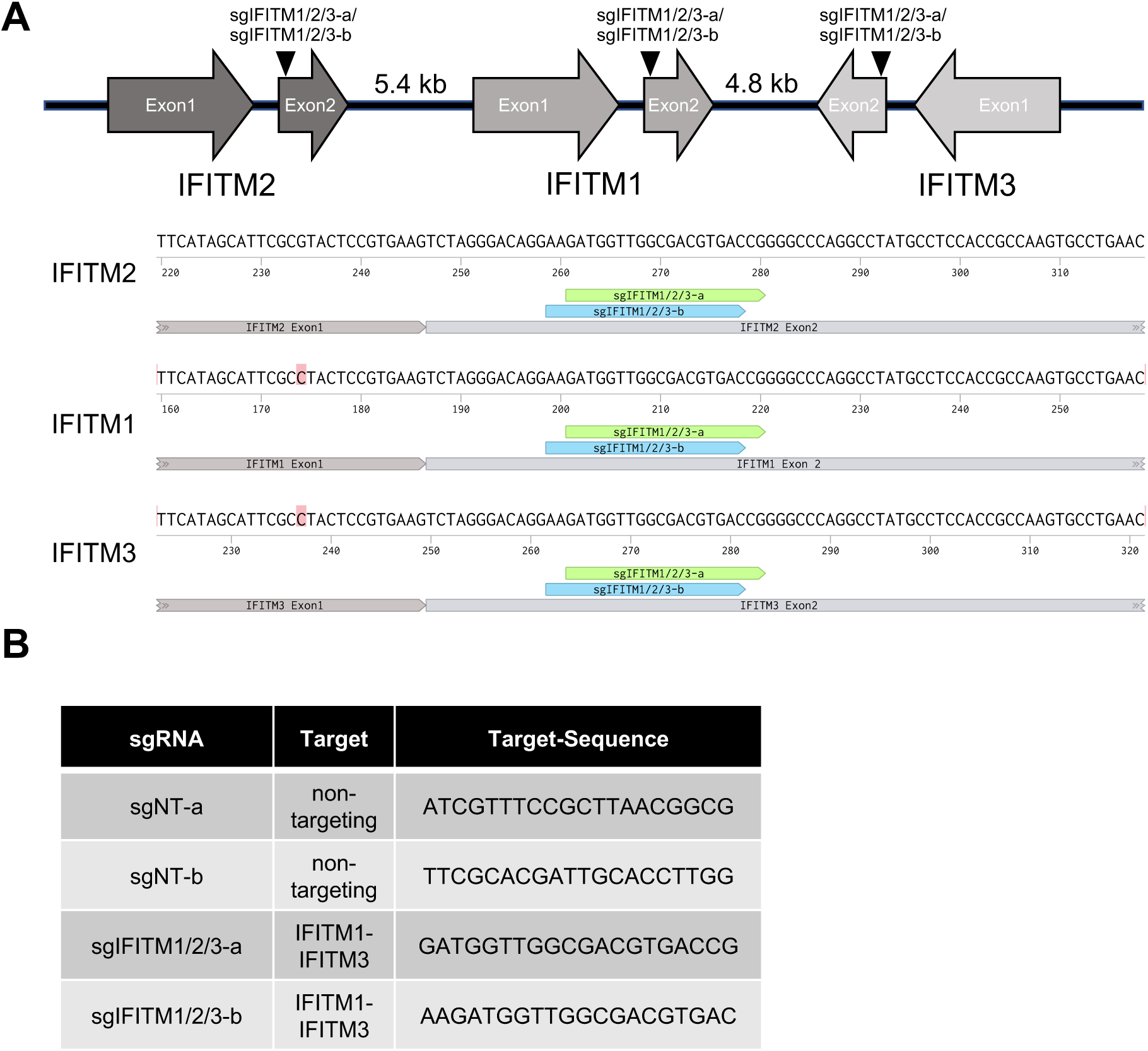
Localization of the IFITM-cluster on chromosome 11 in the human genome and sgRNAs used in this study. **(A)** Schematic drawing (not to scale) of the localization of IFITM1, IFITM2, and IFITM3 on chromosome 11 in the human genome with target sites of sgRNAs targeting exon2 of IFITM1-3 (sgIFITM1/2/3-a, sgIFITM1/2/3-b; upper panel). Alignment of the target sites of sgIFITM1/2/3-a and sgIFITM1/2/3-b (lower panel). **(B)** Sequences of sgRNAs used in this study.

We chose the lung epithelial cell line A549 as an epithelial cell model. KSHV is occasionally detected in lung tissue (58) and A549 are well characterized with regard to IFITM-mediated restriction of different viruses (1, 9, 53). HFF were chosen as a fibroblast model and HUVEC as a model for endothelial cells. Knockout or substantial knockdown of IFITM1, IFITM2, and IFITM3 was achieved (Fig. 3 A-C, right panel). Lentiviral particles (LP) encoding a GFP reporter gene pseudotyped with IAV-hemagglutinin (HA)/neuraminidase (NA) (IAV-LP) served as positive control for IFITM-mediated restriction, while particles pseudotyped with IFITM-resistant amphotropic murine leukemia virus (MLV) envelope (MLV-LP) served as negative control (1). Infections were performed with or without prior IFN-*α* stimulation. IFN-*α* treatment resulted in a significant reduction of KSHV, RRV, and IAV-LP infection in A549 (Fig. 3 A, left panel). Both KSHV and RRV infection were enhanced in non-IFN-*α*-treated IFITM1/2/3 knockout A549, indicating that basal IFITM levels or IFITM expression induced upon contact with the inoculum affect KSHV and RRV infection of A549. In IFN-*α*-treated IFITM1/2/3 knockout cells, infection nearly reached levels of non-IFN-*α*-treated sgNT-transduced cells without IFITM1/2/3 knockout. IAV-LP infection was dramatically increased upon IFITM1/2/3 knockout, while MLV-LP infection was not affected by IFITM1/2/3 knockout in A549, in keeping with published results (1).

**Figure 3.**
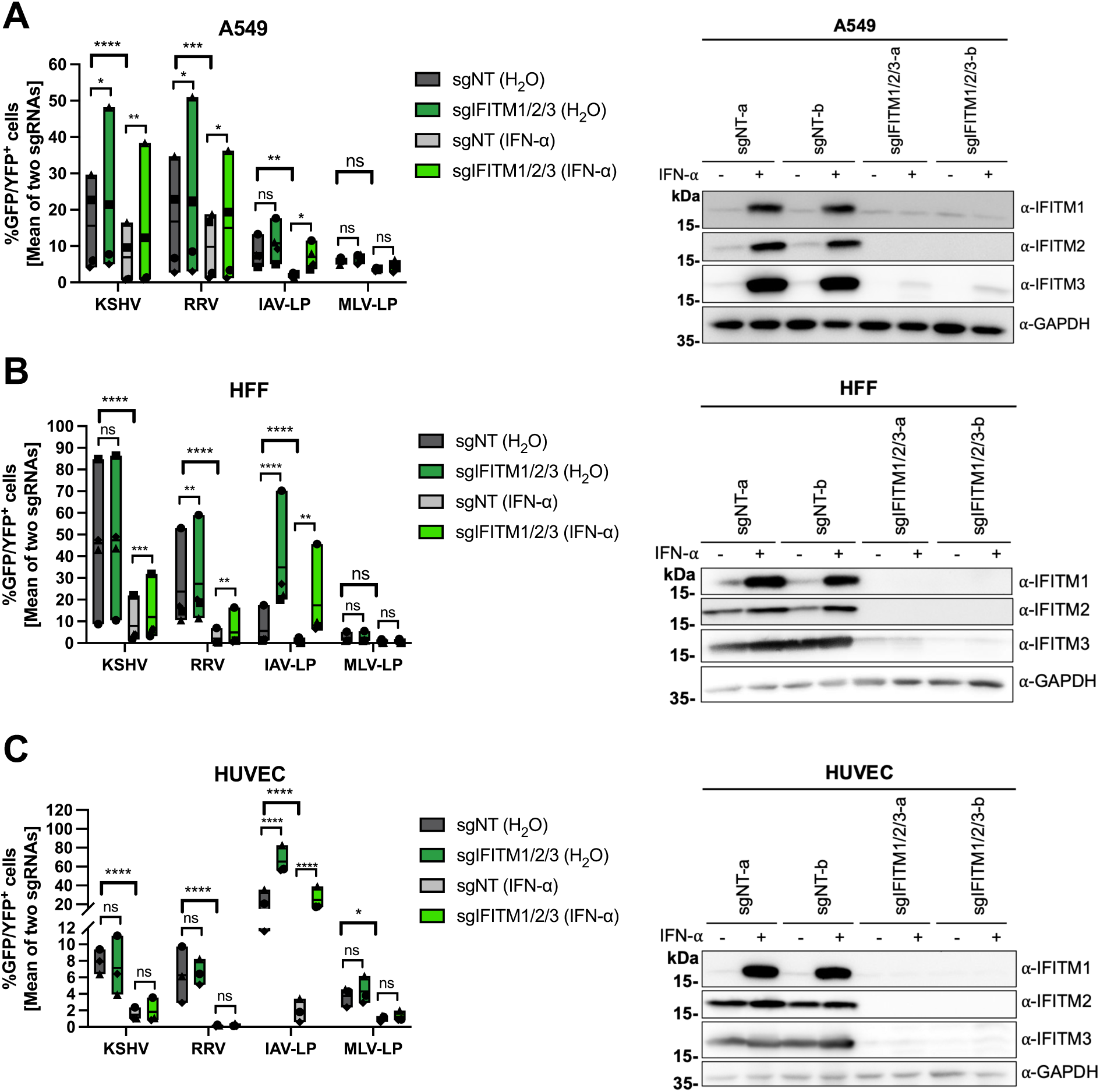
IFITM1/2/3 triple-knockout enhances KSHV and RRV infection in A549 and HFF cells. **(A)** A549, **(B)** HFF and **(C)** HUVEC cells were transduced with lentiviral vectors encoding Cas9 and the sgRNAs shown in Fig. 2. (A-C, left panel) IFITM-knockout (sgIFITM1/2/3-a, sgIFITM1/2/3-b) or control cells (sgNT-a, sgNT-b) treated with IFN-α (5000 U/ml) or H_2_O (control) and infected with KSHV-GFP, RRV-YFP, IAV lentiviral pseudotype (IAV-LP), or MLV lentiviral pseudotype (MLV-LP). Infection was measured using flow cytometry to detect expression of the fluorescent reporter gene. The graph shows individual data points representing averaged values for GFP^+^/YFP^+^ cells of either two non-targeting (sgNT-a, sgNT-b) or IFITM1/2/3 knockout (sgIFITM1/2/3-a, sgIFITM1/2/3-b) transduced cells and floating bars representing the mean averaged from four independent experiments for A549 and HFF (A,B) and three independent experiments for HUVEC (C). Infections for each single experiment were performed in triplicates for each condition. Datapoints from the same experiment are labeled with identical symbols. The different sgRNAs were treated as biological replicates within each experiment. Statistical significance was determined by two-way ANOVA, p-values were corrected for all possible multiple comparisons within one family by Tukey’s method (p>0.05, ns; p≤0.05, *; p≤0.01, **; p≤0.001, ***; p≤0.0001, ****). Representative Western blots (A-C, right panel) of IFITM-knockout (sgIFITM1/2/3-a or sgIFITM1/2/3-b) or control (sgNT-a or sgNT-b) cells treated with IFN-α (5000 U/ml) or H_2_O. Indicated IFITM expression was detected with antibodies shown in Fig. 1A; GAPDH served as loading control.

IFN-*α* pre-treatment reduced KSHV and RRV infection of HFF more potently than infection of A549 (Fig. 3 B, left panel). However, IFITM1/2/3 knockout in HFF only enhanced KSHV infection of IFN-*α*-treated cells, while RRV infection was slightly but significantly enhanced in both IFN-*α* and non-IFN-*α*-treated IFITM1/2/3 knockout cells. We observed relatively high basal IFITM2/3 expression in HFF, which was only marginally increased by IFN-*α* (Fig. 3 B, right panel). Infection of IFN-*α*-treated IFITM1/2/3 knockout HFF by KSHV or RRV did not reach levels of untreated sgNT-transduced cells, unlike what was observed with A549, suggesting that IFITM-mediated restriction of KSHV and RRV infection plays a comparatively minor role in the overall IFN-*α*-mediated restriction of these two herpesviruses in HFF. The most potent effect of IFITM1/2/3 knockout was observed with IAV-LP infection, which was increased in both IFN-*α*- and control-treated cells. MLV-LP infection of HFF was not significantly affected by IFITM1/2/3 knockout. Like HFF, HUVEC expressed IFITM2 and IFITM3 at high basal levels (Fig. 3 C, right panel). IFN-*α* treatment of HUVEC resulted in a reduction of KSHV infection and an even more pronounced reduction of RRV infection (Fig. 3 C, left panel). However, IFITM1/2/3 knockout had no significant effect on KSHV or RRV infection. Again, IAV-LP infection was strongly enhanced by IFITM1/2/3 knockout in both IFN-*α*- and non-treated HUVEC cells, while MLV-LP was not affected.

Overall, these results demonstrate IFITM-mediated restriction of KSHV and RRV infection of A549 and HFF, but not HUVEC.

### IFITM1 overexpression reduces KSHV and RRV infection in a cell type-dependent manner

We next investigated the effect of individual IFITMs through directed expression by retroviral transduction (Fig. 4 A-D, right panel) and included additional cell lines: i) 293T cells as another cell line of either epithelial or neuroendocrine origin (59), and ii) SLK cells, a clear renal carcinoma cell line (60) that is an established model for KSHV infection and propagation (61). We excluded HUVEC here, as IFITM1/2/3 knockout was without effect in these cells, despite high IFITM expression, which makes it unlikely that additional overexpression yields meaningful results.

**Figure 4.**
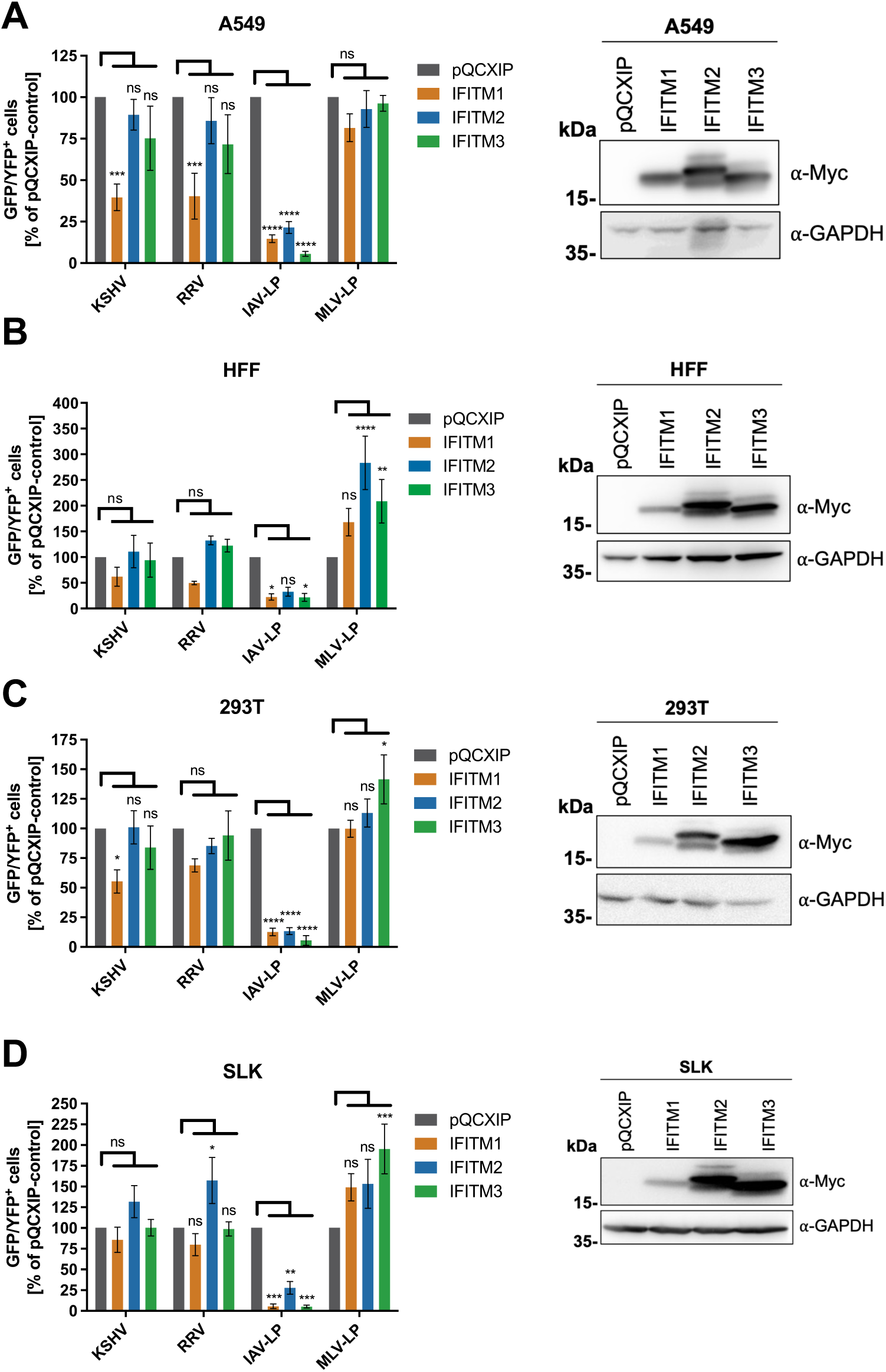
Overexpression of IFITM1 inhibits KSHV and RRV infection in a cell type-dependent manner. **(A)** A549, **(B)** HFF, **(C)** 293T and **(D)** SLK cells were transduced with pQCXIP-constructs to express IFITM1-3 or pQCXIP (empty vector). (A-D, left panel) IFITM overexpressing cells were infected with KSHV-GFP, RRV-YFP, IAV lentiviral pseudotype (IAV-LP) or MLV lentiviral pseudotype (MLV-LP). Infection was measured using flow cytometry to detect expression of the fluorescent reporter genes. The data shows values normalized to pQCXIP empty vector, which was set to 100%, and the error bars represent the standard error of the mean of four independent experiments, each performed in triplicates. Statistical significance was determined by ordinary two-way ANOVA, p-values were corrected for multiple comparisons by Dunnett’s method (p>0.05, ns; p≤0.05, *; p≤0.01, **; p≤0.001, ***; p≤0.0001, ****). Representative Western blots (A-C, right panel) of IFITM-overexpressing cells. Expression of myc-tagged IFITMs was determined using anti-myc antibody; GAPDH served as loading control.

Overexpression of IFITM1 in A549 reduced KSHV and RRV infection by over 50%, whereas overexpression of IFITM2 and IFITM3 only resulted in a non-significant reduction (Figure 4 A left panel), identifying IFITM1 as the IFITM that restricts KSHV and RRV in A549 cells. In agreement with the results in IFITM1/2/3 knockout experiments and published results (1), IAV-LP infection was reduced by all IFITMs, most prominently by IFITM3, and infection of MLV-LP was not affected.

IFITM1 overexpression also reduced RRV and KSHV infection of HFF (Fig. 4 B, left panel), but the effect did not reach statistical significance, mostly because of a rather high pooled variance in this set of experiments, which may reflect the primary nature of HFF combined with comparatively high constitutive IFITM expression. In addition, we observed a non-significant but noticeable enhancement of RRV infection in IFITM2-overexpressing HFF. Again, these observations are in agreement with the effects observed in IFITM1/2/3 knockout HFF. While IAV-LP infection was reduced in HFF, MLV-LP infection was enhanced by overexpression of all IFITMs, significantly for IFITM2 and IFITM3.

A trend similar to that observed in A549 was also observed in 293T (Fig. 4 C, left panel): IFITM1 overexpression reduced KSHV and RRV infection, although not significantly for RRV. MLV-LP infection was slightly increased by overexpression of IFITM3 in 293T.

A different observation was made in SLK (Figure 4 D, left panel), where neither IFITM1 nor IFITM3 overexpression resulted in reduced KSHV or RRV infection. Again, IFITM2 overexpression in SLK slightly enhanced KSHV infection and significantly enhanced RRV infection. An enhancement of infection by all IFITMs was observed with MLV, significantly for IFITM3. Taken together, our IFITM overexpression experiments corroborated the results observed in our IFITM1/2/3 knockout experiments and a cell type-dependent activity of individual IFITMs towards KSHV and RRV. Furthermore, IFITM1 was identified as the major contributor to IFITM-mediated restriction of KSHV and RRV.

### KSHV does not specifically colocalize with IFITMs

IFITM localization was reported to play a critical role in their antiviral effect (62, 63). For the highly restricted IAV, IFITM3-mediated restriction might be partially explained by the observation that IAV specifically colocalizes with IFITM3-positive vesicles (15, 53). As expected, we observed different subcellular localizations of IFITM1, IFITM2, and IFITM3 in IFN-*α*-treated A549 (Fig. 5 A). IFITM2 and IFITM3 partially colocalized with the early endosome marker EEA1 and to a larger extent with the late endosome/lysosome marker LAMP1, while IFITM1 localized to the plasma membrane and was distributed more toward the perimeter of the cell. IFITM1 was also found colocalized with EEA1 and LAMP1, but not as pronounced as IFITM2/IFITM3. Next, we analyzed colocalization of KSHV particles with IFITMs. We utilized a KSHV_mNeon-orf65, which is tagged with mNeonGreen at the capsid protein orf65, to visualize virions in IFN-*α*-treated cells at different timepoints (Fig. 5 B). KSHV_mNeon-orf65 particles were detectable at the perimeter at the 0-min timepoint and were detected inside the cells from the 30-min timepoint on. Some particles reached the nucleus at the 240-min timepoint. As IFITMs are widely distributed throughout the cell, partial overlap with KSHV_mNeon-orf65 particles was observed for all IFITMs, most prominently at later time points and for IFITM1. While some particles localized to areas of high intensity in the IFITM staining, KSHV_mNeon-orf65 particles were also frequently found in regions with overall lower IFITM signal. These areas were often adjacent to IFITM-positive areas, which might be compatible with the luminal spaces of large vesicles.

**Figure 5.**
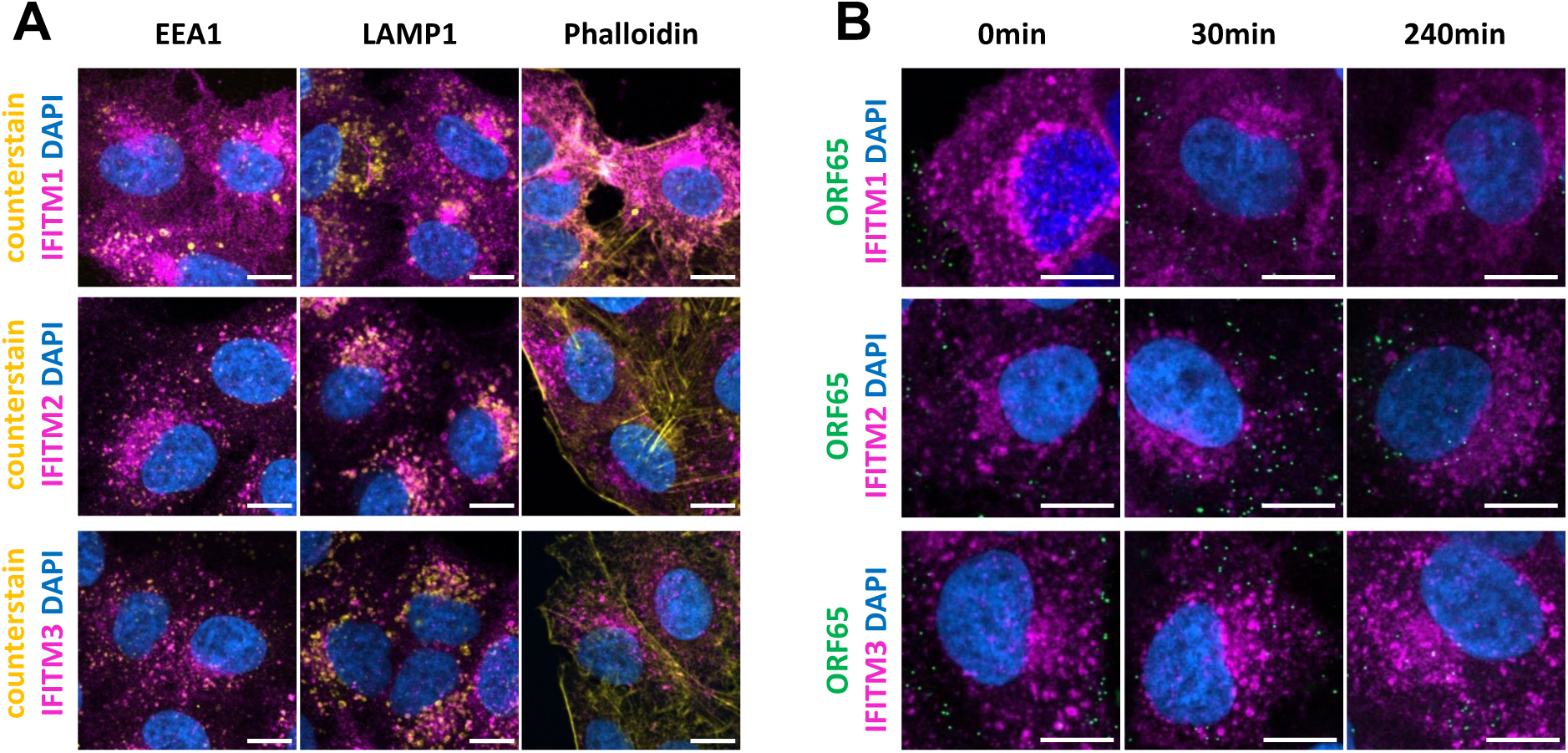
KSHV virus particles do not extensively colocalize with IFITM1, IFITM2, and IFITM3 in A549 cells. **(A)** Confocal microscopy images of IFN-α-treated (5000 U/ml) A549 cells stained with IFITM1, IFITM2, or IFITM3 antibody (magenta). Co-staining was performed with antibodies to EEA1, LAMP1, or phalloidin conjugate (yellow) and Hoechst (blue). **(B)** Confocal microscopy images of IFN-α-treated (5000 U/ml) A549 cells, infected with KSHV_mNeon-orf65 (green). Staining was performed using IFITM1, IFITM2, or IFITM3 antibody (magenta) and Hoechst (blue). The Scale bars represents 10 µm.

### KSHV and RRV glycoprotein-mediated cell-cell fusion is reduced by IFITMs

IFITMs were reported to modulate overall membrane fusogenicity and thereby entry of viral particles (11, 12, 64). We therefore utilized a cell-cell fusion assay to determine whether KSHV and RRV glycoprotein-mediated fusion activity is modulated by IFITMs. 293T effector cells were transfected with KSHV gH/gL or RRV gH/gL together with RRV gB and a plasmid encoding a VP16-Gal4 transactivator fusion protein. RRV gB was used because KSHV gB does not allow for efficient cell-cell fusion (46). Transfected effector cells were added to IFN-*α*-treated A549 IFITM1/2/3 knockout cells transduced with a lentiviral Gal4-driven TurboGFP-luciferase reporter construct or 293T cells transfected to express IFITM1, IFITM2, or IFITM3, and a Gal4-driven TurboGFP-luciferase reporter construct. Luciferase activity was measured as a readout for fusion.

Treatment with IFITM-targeting sgRNAs resulted in an increase of KSHV and RRV gH/gL/gB-mediated cell-cell fusion compared to non-targeting controls (Fig. 6 A). Viral glycoprotein expression and IFITM1/2/3 knockout in target cells was confirmed by Western blot (Fig. 6 B). Under conditions of recombinant overexpression, all three IFITMs were capable of reducing KSHV and RRV gH/gL/gB-mediated cell-cell fusion, with IFITM1 being the most effective (Fig. 6 C). To exclude the possibility that the inhibition of KSHV and RRV glycoprotein-mediated cell-cell fusion occurs in response to changes in cell surface protein composition upon IFITM1/2/3 knockout, we measured cell surface expression of a set of selected cell-surface receptors. Cell surface expression of the KSHV receptors EphA2 and integrin *α*V as well as transferrin receptor (TrfR) remained unchanged upon IFITM1/2/3 knockout (Fig. 6 D). This suggests that IFITMs reduce cell-cell fusion through a mechanism distinct from receptor regulation.

**Figure 6.**
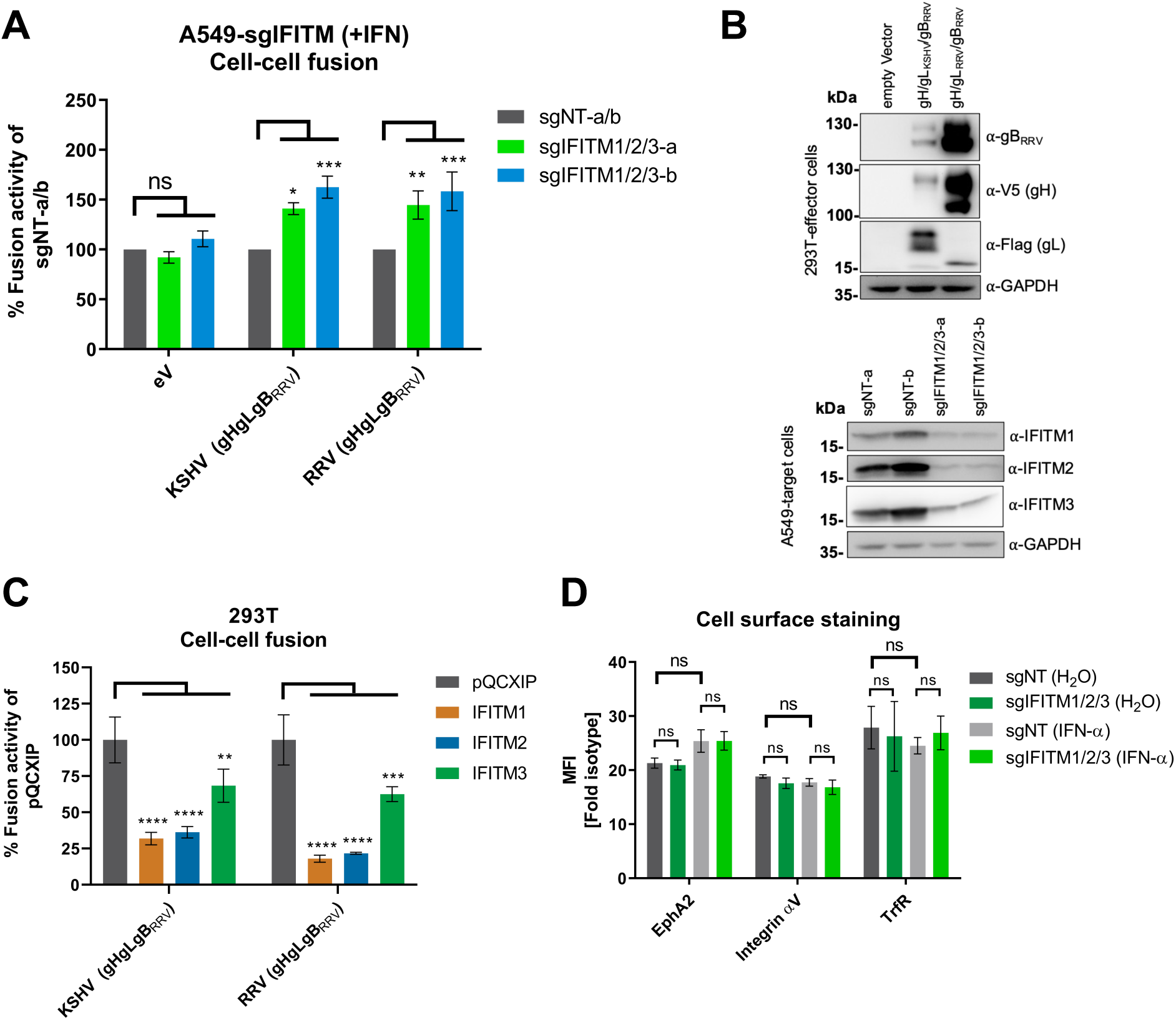
IFITMs inhibit KSHV and RRV glycoprotein-mediated cell-cell fusion. **(A)** Cell-cell fusion assay. Effector cells (293T transfected with either empty vector (eV) or expression plasmids for the indicated viral glycoproteins together with Vp16-Gal4 expression plasmid) were added to target cells (A549 cells transduced with a Gal4-driven TurboGFP-Luciferase construct and the respective CRISPR/Cas9 sgRNA-construct), which had been pre-incubated for 16h with IFN-α (5000 U/ml). After 48 h, luciferase activity was measured. Values were normalized to the mean of the two non-targeting controls sgNT-a and sgNT-b (sgNT-a/b), which was set to 100, for each experiment. Error bars represent standard error of the mean of four independent experiments, each performed in triplicates. Statistical significance was determined by two-way ANOVA; p-values were corrected for multiple comparisons by Dunnet’s method (p>0.05, ns; p≤0.05, *; p≤0.01, **; p≤0.001, ***; p≤0.0001, ****). **(B)** The expression of proteins in 293T effector and A549 target cells after co-cultivation was analyzed by Western blot from lysates harvested for determination of luciferase activity shown in **(A)** using the indicated antibodies. GAPDH served as loading control. **(C)** Cell-cell fusion assay. Effector cells (293T transfected with expression plasmids for the indicated viral glycoproteins together with Vp16-Gal4 expression plasmid) were added to target cells (293T cells transfected with a Gal4-driven TurboGFP-Luciferase construct and the respective pQCXIP-IFITM construct). After 48 h, luciferase activity was measured. Values were averaged from three independent experiments, each performed in triplicates. The data was normalized to empty vector control pQCXIP, which was set to 100, error bars represent the standard deviation. Statistical significance was determined by two-way ANOVA, p-values were corrected for multiple comparisons by Dunnet’s method (p>0.05, ns; p≤0.05, *; p≤0.01, **; p≤0.001, ***; p≤0.0001, ****). **D** A549 cells were transduced with a lentiviral vector encoding Cas9 and sgRNAs shown in Figure 2. IFITM-knockout (sgIFITM1/2/3-a, sgIFITM1/2/3-b) or control cells (sgNT-a and sgNT-b) treated with IFN-α (5000 U/ml) or H_2_O (control) were stained for cell surface expression of indicated proteins. The graph shows values for the mean fluorescence intensity fold over isotype control averaged from two non-targeting (sgNT-a, sgNT-b) or IFITM1/2/3 knockout (sgIFITM1/2/3-a, sgIFITM1/2/3-b) transduced cells from one representative experiment performed in triplicates. Error bars represent the standard deviation. Statistical significance was determined by two-way ANOVA, p-values were corrected for multiple comparisons by Tukey’s method (p>0.05, ns; p≤0.05, *; p≤0.01, **; p≤0.001, ***; p≤0.0001, ****).

## DISCUSSION

Differences in the activity of IFITMs against several members of the herpesvirus family have already been reported (18–21, 23). Here, we report that human IFITM1 inhibits the entry of KSHV and of the closely related rhesus macaque virus RRV in a cell type-dependent manner. We identified inhibition of membrane fusion as a potential mechanism through which IFITMs can modulate KSHV and RRV infection.

Combined knockout of all three IFITMs enabled us to study IFITM-mediated restriction through loss-of-function at expression levels that are induced through IFN signaling and free from potential artefacts through overexpression-induced mislocalization. Our approach revealed that KSHV and RRV infection are enhanced upon IFITM1/2/3 knockout in A549 and HFF but not in HUVEC. This finding may be explained by differences in entry routes that KSHV and RRV utilize to enter these different cell types. KSHV was shown to enter HFF (65) and RRV rhesus fibroblasts (33, 44) via clathrin-mediated endocytosis, whereas KSHV enters HUVEC via macropinocytosis (42). While most viruses that are restricted by IFITMs enter cells via clathrin- or caveolin-mediated endocytosis, only Ebola and Marburg viruses are restricted and enter their target cells predominantly via macropinocytosis (reviewed in (66)). However, Ebola and Marburg virus glycoproteins are activated by endosomal cathepsins, which are mainly found in endolysosomal vesicles that also contain IFITMs (reviewed in (67, 68)). In contrast, KSHV might already fuse in acidified IFITM-negative macropinocytotic compartments and thereby avoid IFITM restriction in HUVEC cells. Although IFITM1/2/3 knockout enhanced KSHV and RRV infection in A549 and HFF, the overall contribution to the IFN-mediated block to infection was different. In HFF, the enhancement was mainly observable in IFN-*α*-treated cells, while in A549 the enhancement was also observable in non-IFN-treated cells. In A549, the IFITM1/2/3 knockout-mediated enhancement practically cancelled out the IFN-*α*-mediated inhibition of KSHV and RRV infection, similar to what was observed for the highly restricted IAV-LP. Unfortunately, the detailed mechanism of KSHV and RRV entry into A549 cells is presently unknown. Differences in the response to IFITM1/2/3 knockout may very well represent differences in the viral entry routes. Despite minor differences, IFITM1/2/3 knockout similarly impacted KSHV and RRV infection, compatible with a broadly acting mechanism like decreasing membrane fusogenicity. Vice versa it was also shown that primate IFITMs are effective against human viruses (69, 70), in line with the high degree of conservation of IFITMs in primate species (71, 72).

Overexpression of individual IFITMs in different cell types revealed IFITM1 as the major contributor to IFITM-mediated restriction of KSHV and RRV. Similar to our observations, an antiviral effect of IFITM1 in A549 cells was also identified for the alphaherpesvirus HSV-1 in IFITM1 overexpression and siRNA-mediated knockdown experiments (18), which suggests broad activity of IFITM1 against herpesviruses. Of note, an effect of IFITM1 on KSHV infection has already been described by Hussein et al.; however, in contrast to our study, their study reported that infection by KSHV, EBV, and HSV-2 was enhanced upon overexpression of IFITM1 in the BJAB B-cell cell line and in HMVEC-D cells (21). While we did not observe this phenomenon in the cell types analyzed in this study, our observations of cell-type dependent antiviral activity do not rule out the possibility that in some cell types infection might actually be enhanced by IFITM1 expression. In line with this notion, overexpression of IFITM2 resulted in a mild enhancement of RRV infection in SLK and HFF (Fig. 4).

IFITM1-mediated inhibition of KSHV or RRV infection was less pronounced than inhibition of IAV glycoprotein-driven entry, and inhibition by IFITM2 and IFITM3 was not observable. This is compatible with our results in colocalization experiments. Several groups reported that IAV colocalized strongly with IFITM3 (15, 53, 73). We were unable to observe a pronounced colocalization of IFITMs with KSHV particles. Rather, KSHV_mNeon-orf65 particles that entered the cell were frequently observed in regions with low IFITM signal. While these findings argue against concentration of IFITMs at viral particles, they would be compatible with indirect mechanisms of action such as rerouting of endocytotic pathways or reduction of membrane fusogenicity.

Mechanistically, we found that IFITMs modulate KSHV and RRV glycoprotein-induced membrane fusion at IFN-*α*-induced levels. Overexpression of IFITM1, IFITM2, and IFITM3 revealed that all three IFITMs can in principle reduce the KSHV and RRV glycoprotein-induced cell-cell fusion to a different degree. It should be noted that overexpression of IFITMs leads to abnormal localization, thereby potentially broadening activity. This supports the theory that all IFITMs are, in principle, capable of restricting fusion (6, 16, 74), which might be counteracted by avoidance of IFITM-positive compartments. In line with our experiments, IFITM overexpression was reported to reduce the fusion activity of other viral fusion proteins including the IAV-HA (12, 13, 15) and severe acute respiratory syndrome coronavirus 2 spike (75) as well as the glycoprotein of the otherwise non-restricted Lassa virus (15). Although cell-cell fusion does not universally mirror virus-cell fusion (76), our findings support a model of IFITM1 rendering the membrane less fusogenic. A general impact of IFITMs on membrane properties is also supported by a report that IFITMs inhibit trophoblast fusion (64). While our approach of a triple knockout was also intended to identify potential synergism between the three IFITMs, it did not do so. In HFF, IFITM1 might even counteract the mild enhancing effect that IFITM2 had on RRV infection (Fig. 4 B). Overexpression of IFITM1 was sufficient to effect inhibition with a similar magnitude as the enhancement that was observed after knockout. In light of our results and a recent report that IFITM3 blocks the IAV fusion process through increasing membrane stiffness (13), one might speculate that the three IFITMs exert their inhibitory activity through a similar mechanism at different locations.

Entry driven by the HA and NA glycoproteins of IAV, a respiratory pathogen, was far more potently restricted by IFITMs in fibroblasts and endothelial cells, particularly at constitutive expression levels, than in A549 lung epithelial cells. KSHV and likely RRV (77) are endotheliotropic viruses and were restricted in lung epithelial cells but not endothelial cells. This suggests that IFITMs, which are constitutively expressed at high levels in HUVEC and fibroblasts, constitute a major line of defense against disseminated infection of extrapulmonary tissues by the respiratory pathogen IAV, and that KSHV and RRV may have evolved to avoid IFITM-mediated restriction in their biological niche.

## MATERIALS AND METHODS

### Cell culture

All cell lines in this study (Table 1) were incubated at 37°C and 5% CO_2_ and cultured in Dulbecco’s modified Eagle medium, high glucose, GlutaMAX, 25 mM HEPES (Thermo Fisher Scientific) supplemented with 10% fetal calf serum (FCS; Thermo Fisher Scientific) and 50 µg/ml gentamicin (PAN-Biotech) (D10) except for HUVEC, which were maintained in standard Endothelial Cell Growth Medium 2 (PromoCell), and iSLK cells, which were maintained in D10 supplemented with 2.5 µg/ml puromycin (InvivoGen) and 250 µg/ml G418 (Carl Roth). IFN-*α* treatment was performed by supplementing the respective culture medium with IFN-*α* 2b (Sigma; 5000 U/ml). For seeding and subculturing of cells, the medium was removed, the cells were washed with phosphate-buffered saline (PBS; PAN-Biotech), and detached with trypsin (PAN-Biotech). All transfections were performed using polyethylenimine (PEI; Polysciences) at a 1:3 ratio (mg DNA/mg PEI) mixed in Opti-MEM (Thermo Fisher Scientific).

**Table 1.** Cell lines.

| Cell line | Origin |
| --- | --- |
| 293T cells | a kind gift from Vladan Rankovic, Göttingen, Germany, and originally purchased from the ATCC |
| A549 | Laboratory of Stefan Pöhlmann, German Primate Center-Leibniz Institute for Primate Research, Göttingen, Germany |
| SLK cells | (RRID:CVCL_9569) (NIH AIDS Research and Reference Reagent program) |
| human foreskin fibroblasts (HFF) | (laboratory of Klaus Korn, Universitätsklinikum Erlangen, Institute for Clinical and Molecular Virology, Erlangen, Germany) |
| rhesus monkey fibroblasts (RF) | (laboratory of Prof. Rüdiger Behr, German Primate Center-Leibniz Institute for Primate Research, Göttingen, Germany) |
| Human vascular endothelial cells (HUVEC) | (PromoCell) |
| iSLK cells | A kind gift from Don Ganem (61) |

### Retroviral vectors and pseudotyped lentiviral particles

Retroviruses, lentiviruses and lentiviral pseudotypes were produced by PEI-mediated transfection of 293T cells (see Table 2 for plasmids). For retrovirus production, plasmids encoding gag/pol, pMD2.G encoding VSV-G, and the respective pQCXIP-contructs were transfected (ratio 1.6:1:1.6). For production of lentiviruses used for transduction, psPAX2 encoding gag/pol, pMD2.G encoding VSV-G and the respective lentiviral construct, Gal4-driven TurboGFP-luciferase reporter lentivirus (AX526) or plentiCRISPRv2 were used (ratio 2.57:1:3.57). For lentiviral pseudotypes psPAX2, pLenti CMV GFP Neo and expression plasmids for pCAGGS IAV_WSN-HA and pCAGGS IAV_WSN-NA for IAV-LP or paMLV_env for MLV-LP were used (ratio 1:1.4:2.4). Viruses were harvested twice, 24-48 h and 72-96 h after transfection, passed through a 0.45-µm CA filter and frozen at -80°C. Transduction was performed by adding retroviruses and lentiviruses to cells for 48 h. Afterwards, selection was performed using 10 µg/mL puromycin (InvivoGen; pQCXIP and plentiCRISPRv2 constructs) or 10 µg/mL blasticidin (InvivoGen; AX526 lentivirus).

**Table 2.**
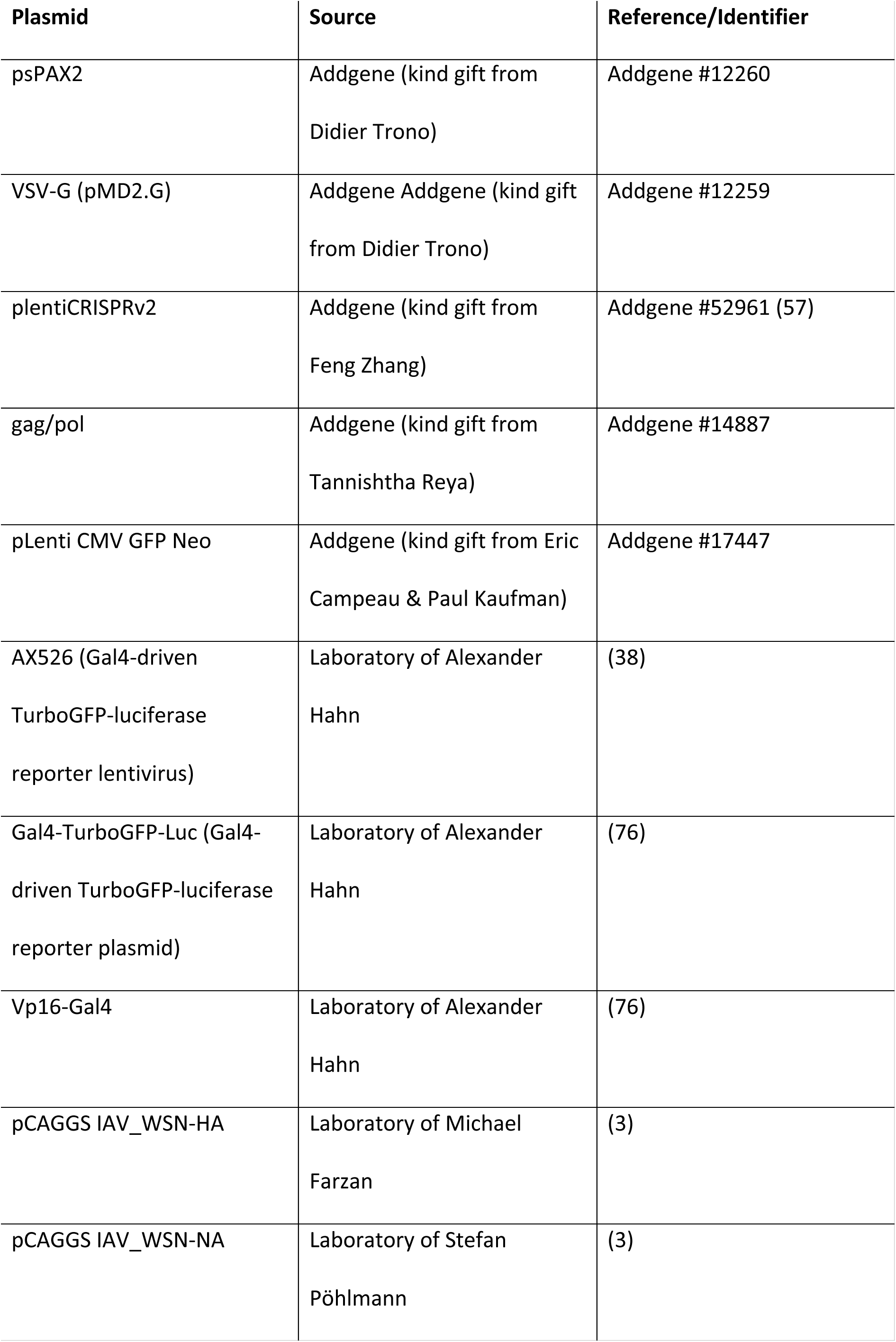
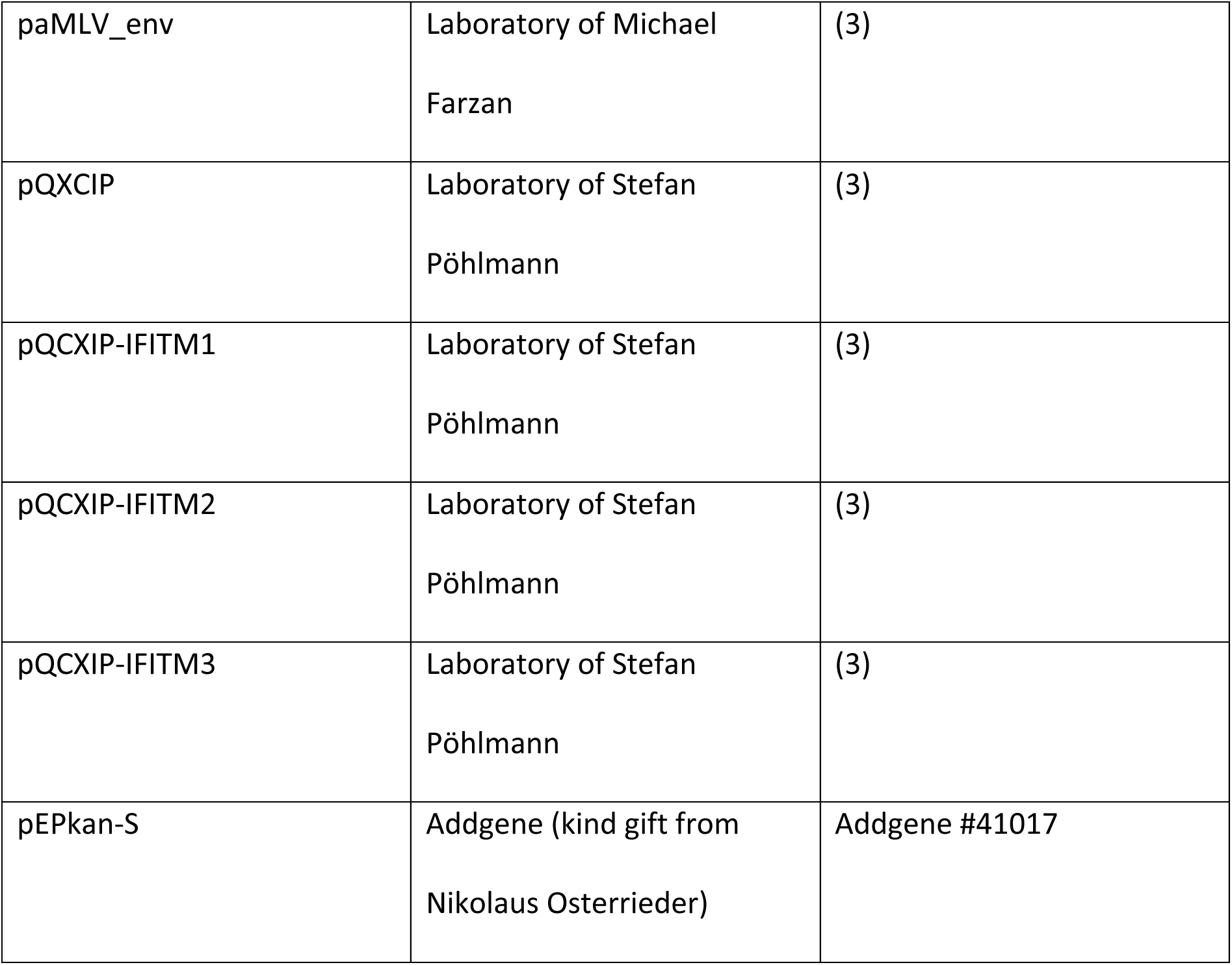
Plasmids.

| Plasmid | Source | Reference/Identifier |
| --- | --- | --- |
| psPAX2 | Addgene (kind gift from<br>Didier Trono) | Addgene #12260 |
| VSV-G (pMD2.G) | Addgene Addgene (kind gift<br>from Didier Trono) | Addgene #12259 |
| plentiCRISPRv2 | Addgene (kind gift from<br>Feng Zhang) | Addgene #52961 (57) |
| gag/pol | Addgene (kind gift from<br>Tannishtha Reya) | Addgene #14887 |
| pLenti CMV GFP Neo | Addgene (kind gift from Eric<br>Campeau & Paul Kaufman) | Addgene #17447 |
| AX526 (Gal4-driven<br>TurboGFP-luciferase<br>reporter lentivirus) | Laboratory of Alexander<br>Hahn | (38) |
| Gal4-TurboGFP-Luc (Gal4-driven TurboGFP-luciferase<br>reporter plasmid) | Laboratory of Alexander<br>Hahn | (76) |
| Vp16-Gal4 | Laboratory of Alexander<br>Hahn | (76) |
| pCAGGS IAV_WSN-HA | Laboratory of Michael<br>Farzan | (3) |
| pCAGGS IAV_WSN-NA | Laboratory of Stefan<br>Pöhlmann | (3) |
| paMLV_env | Laboratory of Michael<br>Farzan | (3) |
| pQXCIP | Laboratory of Stefan<br>Pöhlmann | (3) |
| pQCXIP-IFITM1 | Laboratory of Stefan<br>Pöhlmann | (3) |
| pQCXIP-IFITM2 | Laboratory of Stefan<br>Pöhlmann | (3) |
| pQCXIP-IFITM3 | Laboratory of Stefan<br>Pöhlmann | (3) |
| pEPkan-S | Addgene (kind gift from<br>Nikolaus Osterrieder) | Addgene #41017 |

### Production of KSHV, KSHV_mNeon-orf65, and RRV

For the construction of KSHV_mNeon-orf65, the GFP open reading frame of BAC16 was replaced with a Zeocin resistance gene by amplifying the resistance gene from pcDNA6 (Invitrogen) using Phusion PCR (NEB) and primers BAC16_downstream_of_GFP_STOP_overhang_plus_Zeo_3ʹ and BAC16_upstream_of_GFP_ATG_antisense_strand_overhang_plus_EM7_P_start and inserting it into BAC16 via recombination. A shuttle construct, Ax185_ pCNSmNeonGreen_Kana, was created by inserting the i-SceI/Kanamycin cassette of pEPkan-S (78) into pNCSmNeonGreen using primers mNeonGreen_463-482_for plus mNeonGreen_504-523_rev for the vector and EPKansS_reverse_mNeon_463-482_ov plus EPKans_forward_mNeon_504-523_ov for the insert, followed by Gibson assembly. KSHV_mNeon-orf65 was generated by inserting the mNeonGreen cassette 5ʹ of the first amino acid of orf65 with the addition of a glycine-serine linker according to the protocol described by Tischer et al. (78). The recombination cassette was generated using primers mNeon-GS-KSHVorf65_for plus mNeon-GS-KSHVorf65_rev and Ax185_ pCNSmNeonGreen_Kana as a template. Infectious KSHV and RRV reporter viruses were produced as described previously (36). See Table 3 for oligonucleotide sequences.

**Table 3.**
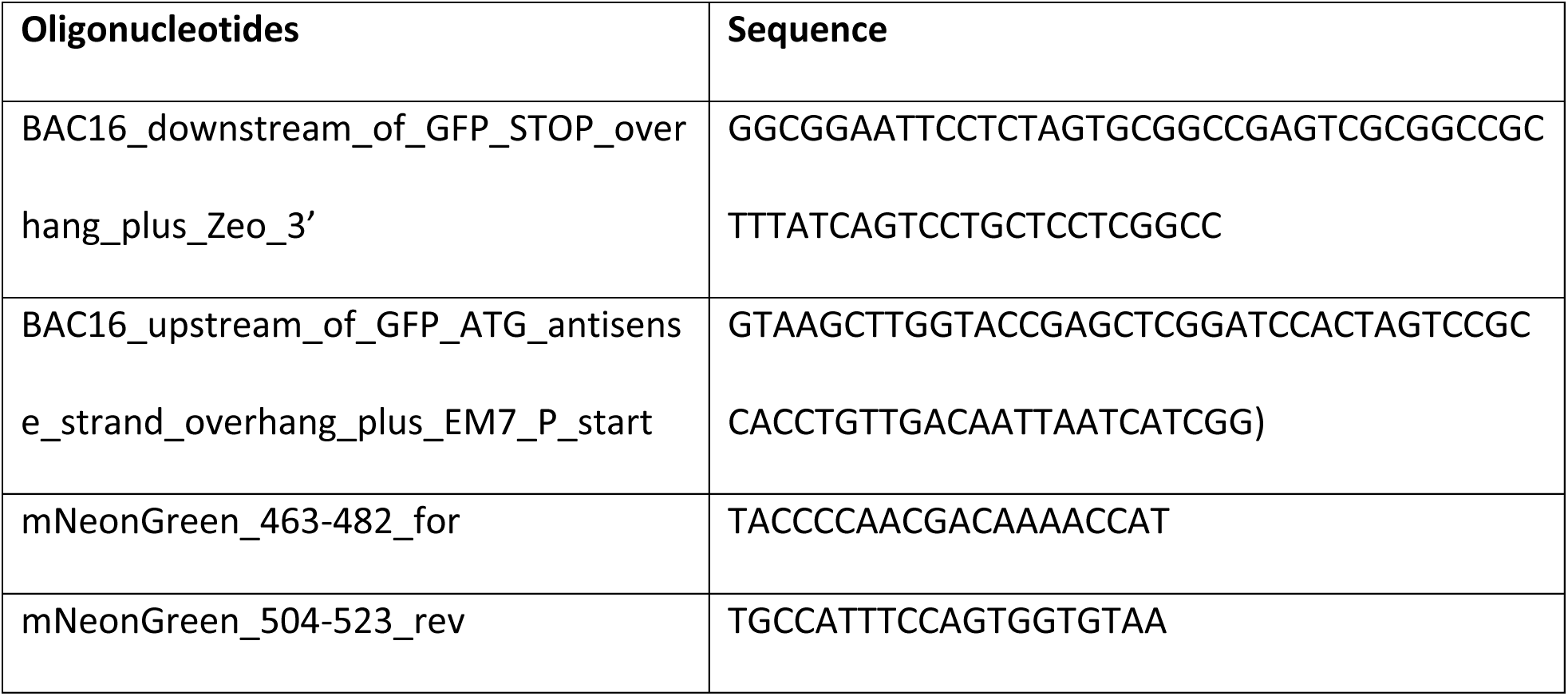
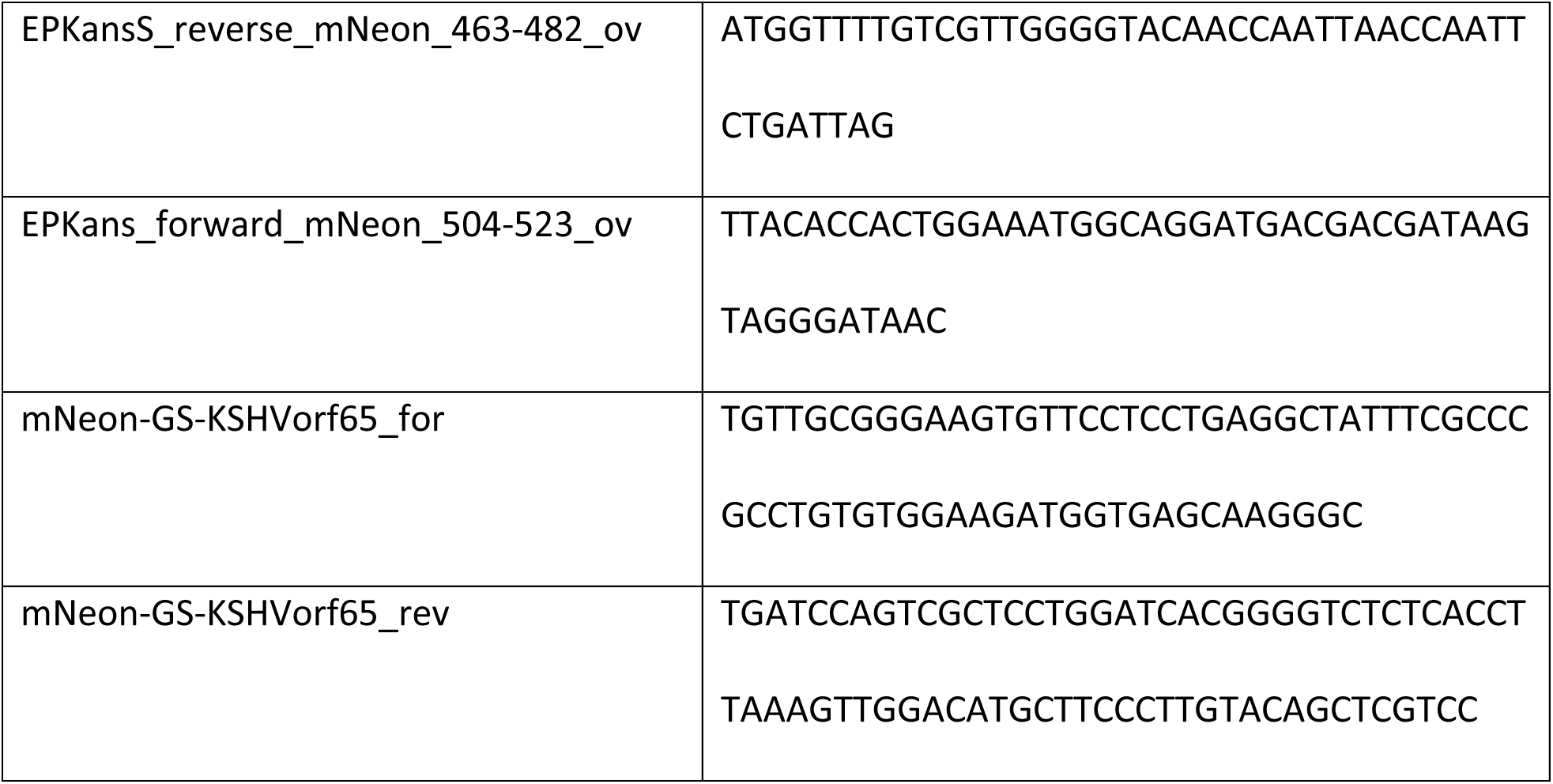
Oligonucleotides.

### Western blot

Western blotting was performed as described previously (36) using the respective antibodies (Table 4).

**Table 4.**
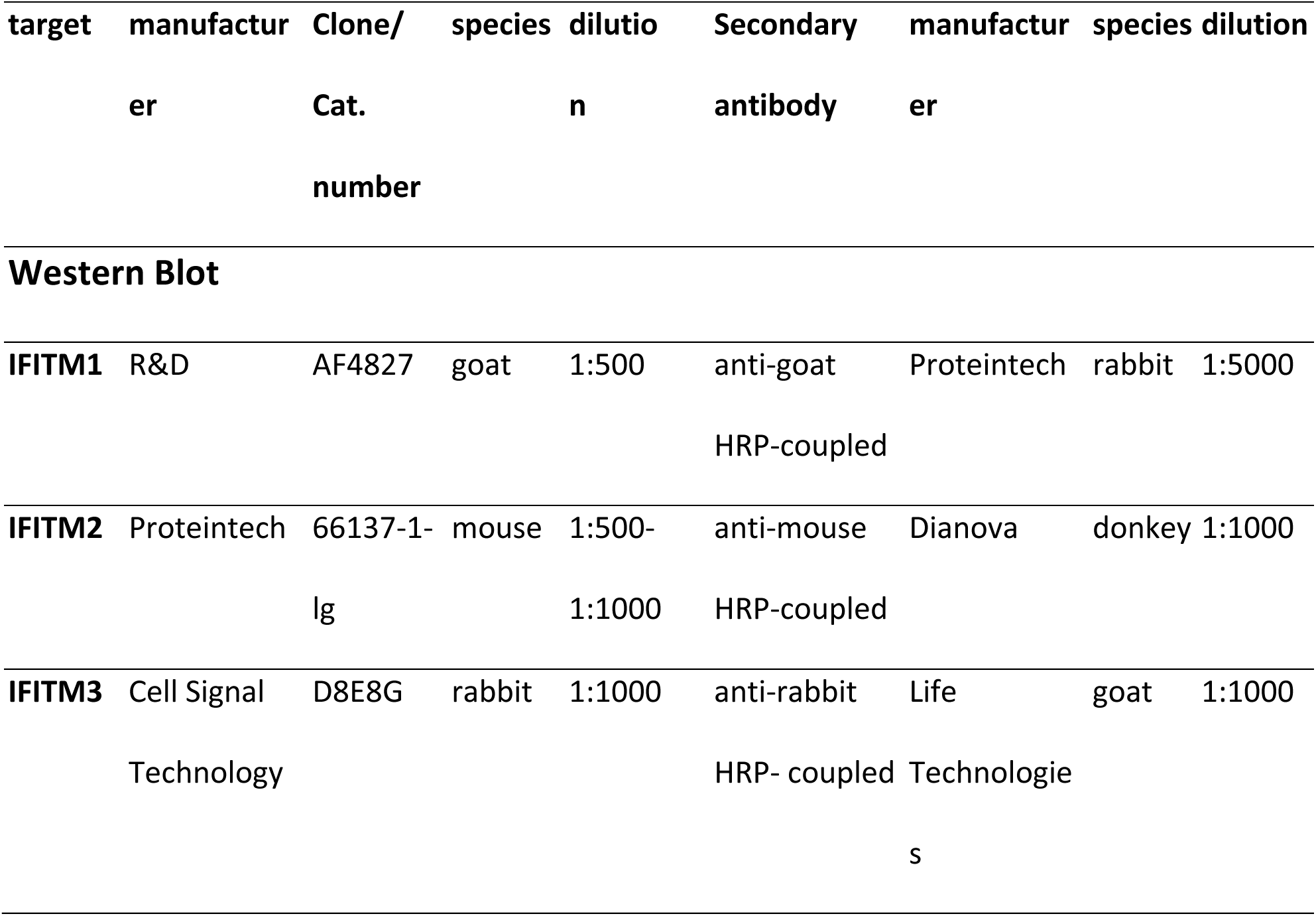
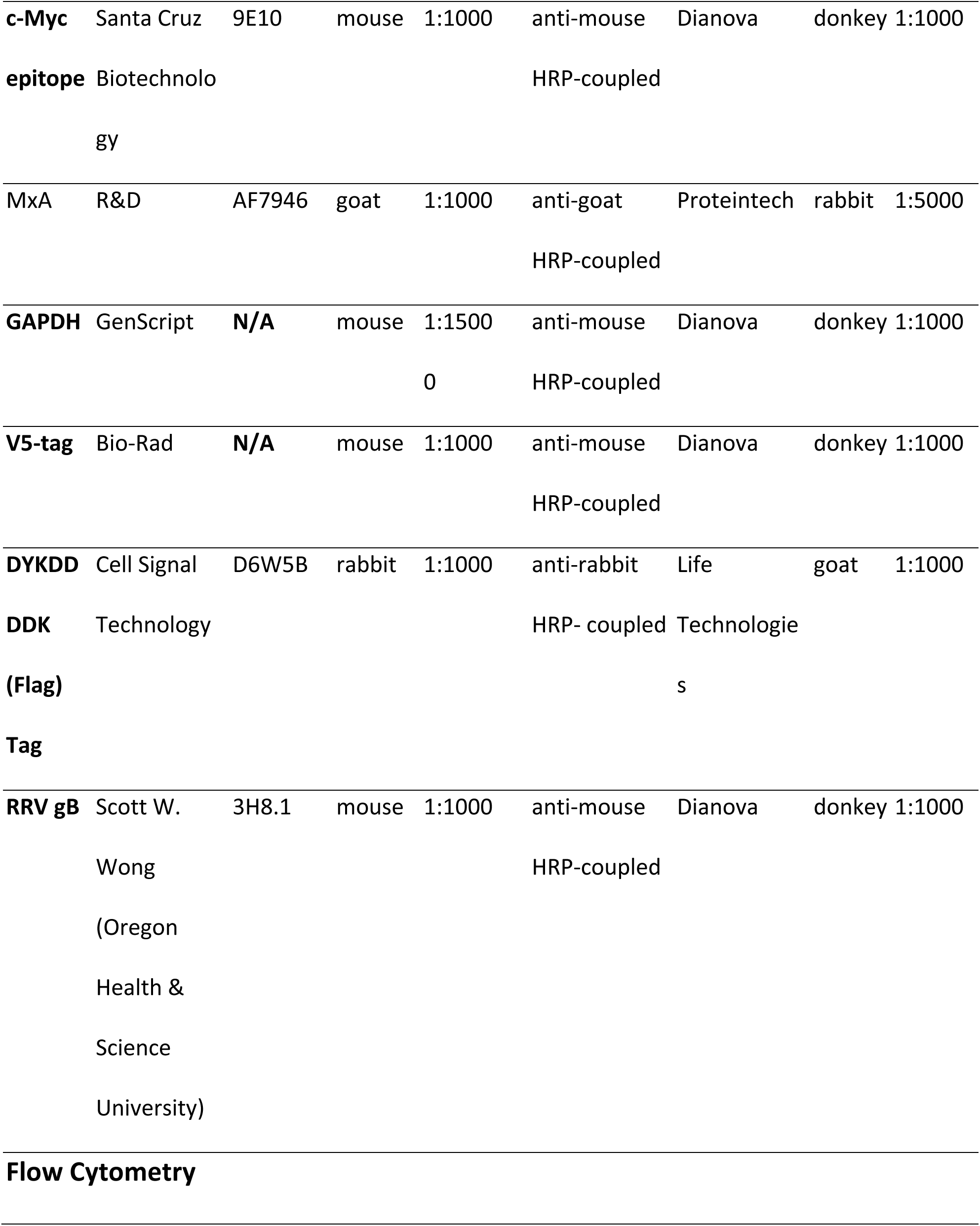
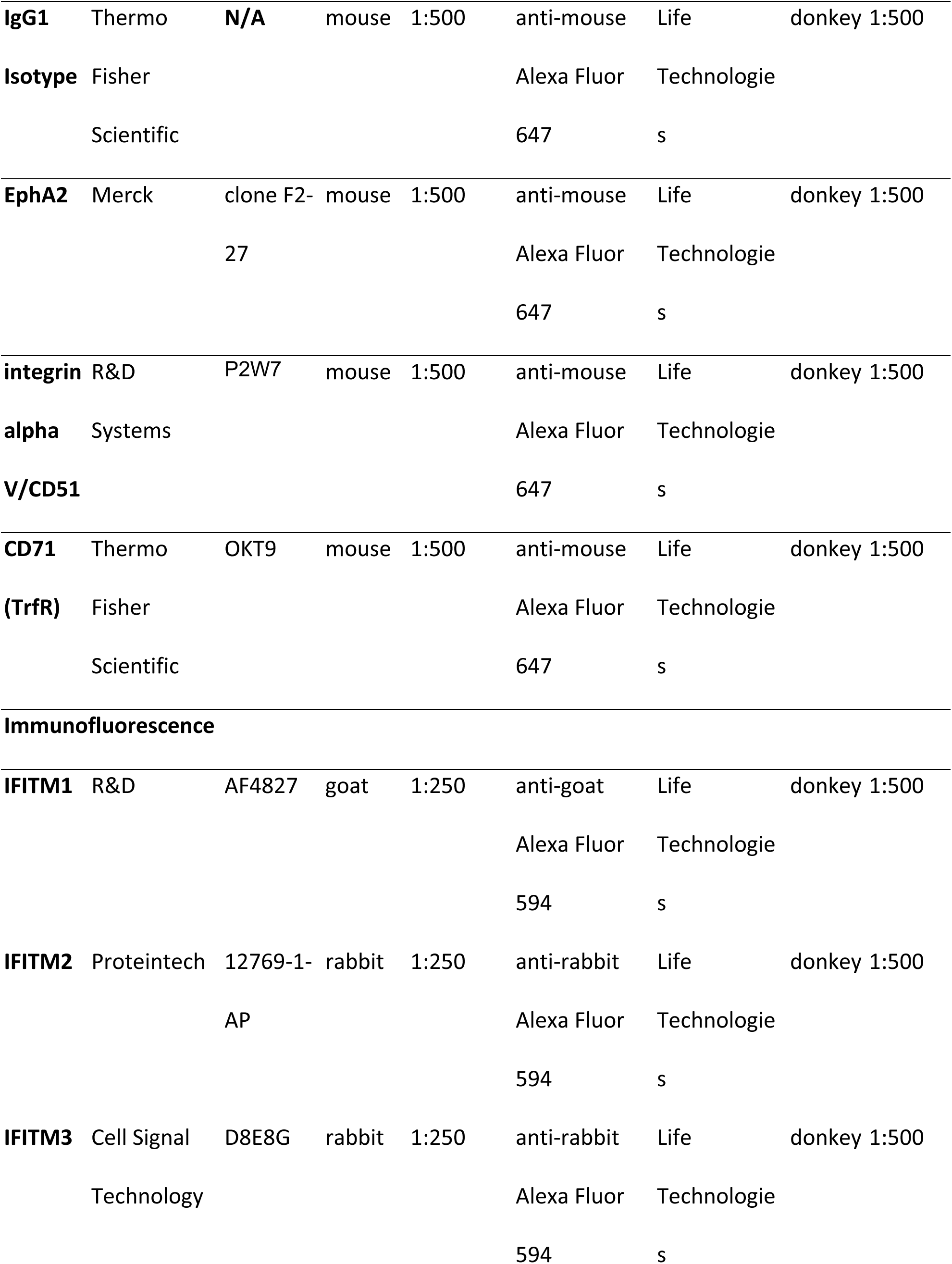
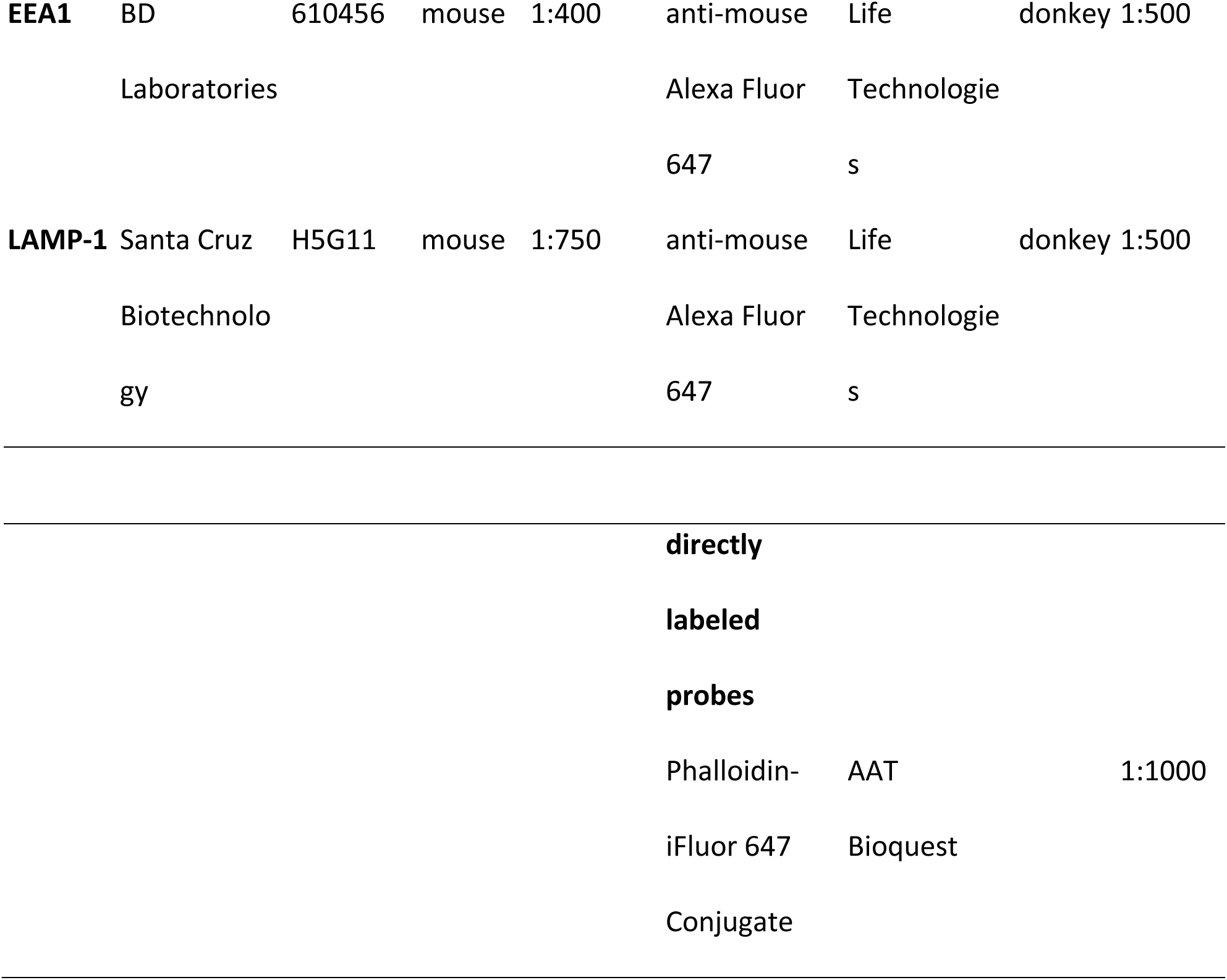
Antibodies.

### CRISPR/Cas9-mediated knockout of immune-related IFITMs

IFITM1-, IFITM2-, and IFITM3-knockout cell pools were generated by CRISPR/Cas9-mediated knockout following the protocol described by Sanjana et al. (57), except that PEI transfection was used. In short, the cells intended for knockout were transduced with lentiviruses harboring the CRISPR/Cas9 gene and sgRNAs targeting IFITM1-3 (sgIFITM1/2/3-a, sgIFITM1/2/3-b) or non-targeting sgRNAs (sgNT-a, sgNT-b). For detection of CRISPR/Cas9-mediated knockout, the cells were treated with IFN-*α* (5000 U/ml) for 16 h. Thereafter, the cells were harvested and subjected to Western blot analysis.

### Infection experiments

IFITM-overexpressing cells were seeded in 48-well plates at 90% confluency 16 h prior to infection. IFITM1/2/3 knockout cells were seeded in 48-well-plates at 70%-80% confluency. After attachment, cells were treated with IFN-*α* (5000 U/ml) or H_2_O (control) for 16 h prior to infection with either KSHV, RRV, IAV-LP or MLV-LP. 48 h post infection, cells were trypsinized, trypsin activity was inhibited by adding 5% FCS in PBS, and the cells were washed and fixed with 4% methanol-free formaldehyde (Roth) in PBS. Infection was determined by detection of GFP^+^/YFP^+^ cells using a LSRII flow cytometer; at least 5000 cells were analyzed.

### Cell-cell fusion assay

293T effector cells were seeded in 6-well plates or 10-cm dishes at 70%-80% confluency and transfected with either empty vector, gH/gL_KSHV_ gB_RRV_ or gH/gL_RRV_gB_RRV_, and Vp16-Gal4. 293T transfected with Gal4-TurboGFP-Luc and pQCXIP-IFITM1-3 were seeded in 48-well plates at 50,000 cells/well. A549 double transduced with lentiviruses encoding a Gal4-driven TurboGFP-luciferase reporter and lentiviruses encoding the CRISPR/Cas9 gene and the respective sgRNAs were seeded in 96-well-plates at 20,000 cells/well, 6 h after seeding the cells were treated with IFN-*α* (5000 U/ml) for 16 h. Cell-cell fusion was started by adding the glycoprotein-expressing effector cells to the target cells in a 1:1 ratio. After 48 h, the cells were lysed in Luciferase Cell Culture Lysis Reagent (Promega) and luciferase activity was determined using Beetle-Juice luciferase assay (PJK Biotech) according to the manufacturer’s instructions and a BioTek Synergy 2 plate reader.

### Flow cytometry

For detection of cell surface proteins, A549 IFITM1/2/3 knockout cells were H_2_O or IFN-*α* treated for 16 h, washed with PBS, detached using EDTA/EGTA (5 mM/5 mM) at 37°C and washed with cold PBS (4°C). The cells were fixed with 4% methanol-free formaldehyde for 5min and washed twice with PBS. Following blocking with 10% FCS (blocking buffer) in PBS, the cells were incubated with primary antibody (Table 4) in blocking buffer for 90 min at 4°C. After washing with PBS, the cells were incubated with secondary antibody (Table 4) in blocking buffer for 45 min at RT in the dark. The cells were washed and post-fixed with 2% methanol-free formaldehyde in PBS. Analysis was performed using an LSRII flow cytometer (BD Biosciences) and Flowing software (University of Turku, version 2.5).

### Immunofluorescence

A549 were seeded on 12-mm coverslips (YX03.1, Carl Roth) in 24-well plates at 150,000 cells/well. After attachment, the cells were treated with either H_2_O (control) or IFN-*α* (5000 U/ml) for 16 h. After 24 h, cold KSHV_mNEON-ORF65 was added. Cells were centrifuged (4,200 rpm, 4°C, 30 min), followed by a 10-min incubation at 4°C. After 3 washes with cold PBS, cells were either fixed in 4% methanol-free formaldehyde in PBS for 10 min (0-min timepoint) or shifted to 37°C after addition of D10. At the indicated timepoints, cells were washed once in PBS and fixed in 4% methanol-free formaldehyde in PBS for 10 min. After fixation, cells were washed three times in PBS. Cell permeabilization and blocking was performed in IF buffer (5% FCS, 0.05% saponin (Sigma) in PBS) for 1 h. Primary antibody (see Table 4) incubation was performed in IF buffer overnight at 4°C. Secondary antibody (see Table 4) incubation or incubation with a directly labeled phalloidin probe was performed after three washes with IF buffer for 1 h at RT. Cells were washed once in IF buffer and stained with Hoechst 33342 1:10000 in PBS (#62249, Thermo Scientific) for 5 min, followed by a final wash with PBS. The coverslips were dried and mounted in anti-Fade Fluorescence Mounting Medium (ab104135, abcam). Images were acquired on a confocal laser scanning microscope (Zeiss LSM800). Laser intensity and signal amplification were maintained between different conditions for each antibody staining. All images were processed using Fiji/ImageJ software.

## ACKNOWLEDGEMENTS

This work was supported by grants from the Deutsche Forschungsgemeinschaft (www.dfg.de, HA 6013/4-1) and the Wilhelm-Sander Foundation (www.wilhelmsander-stiftung.de, project 2019.027.1).

We thank Stefan Pöhlmann and Michael Farzan for sharing plasmids; Klaus Korn for sharing human foreskin fibroblasts; Rüdiger Behr for sharing rhesus monkey fibroblasts; Scott Wong for sharing RRV gB antibodies.

## REFERENCES

1. Brass AL, Huang I-C, Benita Y, John SP, Krishnan MN, Feeley EM, Ryan BJ, Weyer JL, van der Weyden L, Fikrig E, Adams DJ, Xavier RJ, Farzan M, Elledge SJ. 2009. The IFITM Proteins Mediate Cellular Resistance to Influenza A H1N1 Virus, West Nile Virus, and Dengue Virus. 7. Cell 139:1243–1254.

2. Huang I-C, Bailey CC, Weyer JL, Radoshitzky SR, Becker MM, Chiang JJ, Brass AL, Ahmed AA, Chi X, Dong L, Longobardi LE, Boltz D, Kuhn JH, Elledge SJ, Bavari S, Denison MR, Choe H, Farzan M. 2011. Distinct Patterns of IFITM-Mediated Restriction of Filoviruses, SARS Coronavirus, and Influenza A Virus. 1. PLoS Pathog 7:e1001258.

3. Wrensch F, Karsten CB, Gnirß K, Hoffmann M, Lu K, Takada A, Winkler M, Simmons G, Pöhlmann S. 2015. Interferon-Induced Transmembrane Protein–Mediated Inhibition of Host Cell Entry of Ebolaviruses. suppl 2. J Infect Dis 212:S210–S218.

4. Shi G, Kenney AD, Kudryashova E, Zani A, Zhang L, Lai KK, Hall-Stoodley L, Robinson RT, Kudryashov DS, Compton AA, Yount JS. 2021. Opposing activities of IFITM proteins in SARS-CoV-2 infection. EMBO J 40.

5. Mudhasani R, Tran JP, Retterer C, Radoshitzky SR, Kota KP, Altamura LA, Smith JM, Packard BZ, Kuhn JH, Costantino J, Garrison AR, Schmaljohn CS, Huang I-C, Farzan M, Bavari S. 2013. IFITM-2 and IFITM-3 but Not IFITM-1 Restrict Rift Valley Fever Virus. Journal of Virology 87:8451–8464.

6. Bailey CC, Zhong G, Huang I-C, Farzan M. 2014. IFITM-Family Proteins: The Cell’s First Line of Antiviral Defense. 1. Annu Rev Virol 1:261–283.

7. Siegrist F, Ebeling M, Certa U. 2011. The Small Interferon-Induced Transmembrane Genes and Proteins. Journal of Interferon & Cytokine Research 31:183–197.

8. Weston S, Czieso S, White IJ, Smith SE, Kellam P, Marsh M. 2014. A Membrane Topology Model for Human Interferon Inducible Transmembrane Protein 1. 8. PLoS ONE 9:e104341.

9. Feeley EM, Sims JS, John SP, Chin CR, Pertel T, Chen L-M, Gaiha GD, Ryan BJ, Donis RO, Elledge SJ, Brass AL. 2011. IFITM3 Inhibits Influenza A Virus Infection by Preventing Cytosolic Entry. 10. PLoS Pathog 7:e1002337.

10. Narayana SK, Helbig KJ, McCartney EM, Eyre NS, Bull RA, Eltahla A, Lloyd AR, Beard MR. 2015. The Interferon-induced Transmembrane Proteins, IFITM1, IFITM2, and IFITM3 Inhibit Hepatitis C Virus Entry. 43. J Biol Chem 290:25946–25959.

11. Li K, Markosyan RM, Zheng Y-M, Golfetto O, Bungart B, Li M, Ding S, He Y, Liang C, Lee JC, Gratton E, Cohen FS, Liu S-L. 2013. IFITM Proteins Restrict Viral Membrane Hemifusion. PLOS Pathogens 9:18.

12. Desai TM, Marin M, Chin CR, Savidis G, Brass AL, Melikyan GB. 2014. IFITM3 Restricts Influenza A Virus Entry by Blocking the Formation of Fusion Pores following Virus-Endosome Hemifusion. 4. PLoS Pathog 10:e1004048.

13. Guo X, Steinkühler J, Marin M, Li X, Lu W, Dimova R, Melikyan GB. 2021. Interferon-Induced Transmembrane Protein 3 Blocks Fusion of Diverse Enveloped Viruses by Altering Mechanical Properties of Cell Membranes. ACS Nano 15:8155–8170.

14. Amini-Bavil-Olyaee S, Choi YJ, Lee JH, Shi M, Huang I-C, Farzan M, Jung JU. 2013. The Antiviral Effector IFITM3 Disrupts Intracellular Cholesterol Homeostasis to Block Viral Entry. Cell Host Microbe 13:452–464.

15. Suddala KC, Lee CC, Meraner P, Marin M, Markosyan RM, Desai TM, Cohen FS, Brass AL, Melikyan GB. 2019. Interferon-induced transmembrane protein 3 blocks fusion of sensitive but not resistant viruses by partitioning into virus-carrying endosomes. 1. PLoS Pathog 15:e1007532.

16. Shi G, Schwartz O, Compton AA. 2017. More than meets the I: the diverse antiviral and cellular functions of interferon-induced transmembrane proteins. 1. Retrovirology 14:53.

17. Li C, Du S, Tian M, Wang Y, Bai J, Tan P, Liu W, Yin R, Wang M, Jiang Y, Li Y, Zhu N, Zhu Y, Li T, Wu S, Jin N, He F. 2018. The Host Restriction Factor Interferon-Inducible Transmembrane Protein 3 Inhibits Vaccinia Virus Infection. Front Immunol 9:228.

18. Smith SE, Busse DC, Binter S, Weston S, Diaz Soria C, Laksono BM, Clare S, Van Nieuwkoop S, Van den Hoogen BG, Clement M, Marsden M, Humphreys IR, Marsh M, de Swart RL, Wash RS, Tregoning JS, Kellam P. 2018. Interferon-Induced Transmembrane Protein 1 Restricts Replication of Viruses That Enter Cells via the Plasma Membrane. J Virol 93:e02003–18, /jvi/93/6/JVI.02003-18.atom.

19. Warren CJ, Griffin LM, Little AS, Huang I-C, Farzan M, Pyeon D. 2014. The Antiviral Restriction Factors IFITM1, 2 and 3 Do Not Inhibit Infection of Human Papillomavirus, Cytomegalovirus and Adenovirus. 5. PLoS ONE 9:e96579.

20. Xie M, Xuan B, Shan J, Pan D, Sun Y, Shan Z, Zhang J, Yu D, Li B, Qian Z. 2015. Human Cytomegalovirus Exploits Interferon-Induced Transmembrane Proteins To Facilitate Morphogenesis of the Virion Assembly Compartment. 6. J Virol 89:3049–3061.

21. Hussein HAM, Akula SM. 2017. miRNA-36 inhibits KSHV, EBV, HSV-2 infection of cells via stifling expression of interferon induced transmembrane protein 1 (IFITM1). 1. Sci Rep 7:17972.

22. Hussein HAM, Briestenska K, Mistrikova J, Akula SM. 2018. IFITM1 expression is crucial to gammaherpesvirus infection, in vivo. 1. Sci Rep 8:14105.

23. Tartour K, Nguyen X-N, Appourchaux R, Assil S, Barateau V, Bloyet L-M, Burlaud Gaillard J, Confort M-P, Escudero-Perez B, Gruffat H, Hong SS, Moroso M, Reynard O, Reynard S, Decembre E, Ftaich N, Rossi A, Wu N, Arnaud F, Baize S, Dreux M, Gerlier D, Paranhos-Baccala G, Volchkov V, Roingeard P, Cimarelli A. 2017. Interference with the production of infectious viral particles and bimodal inhibition of replication are broadly conserved antiviral properties of IFITMs. PLoS Pathog 13:e1006610.

24. Damania B, Desrosiers RC. 2001. Simian homologues of human herpesvirus 8. Phil Trans R Soc Lond B 356:535–543.

25. Wen KW, Damania B. 2010. Kaposi sarcoma-associated herpesvirus (KSHV): Molecular biology and oncogenesis. Cancer Letters 289:140–150.

26. Chen Q, Chen J, Li Y, Liu D, Zeng Y, Tian Z, Yunus A, Yang Y, Lu J, Song X, Yuan Y. Kaposi’s sarcoma herpesvirus is associated with osteosarcoma in Xinjiang populations 8.

27. Polizzotto MN, Uldrick TS, Hu D, Yarchoan R. 2012. Clinical Manifestations of Kaposi Sarcoma Herpesvirus Lytic Activation: Multicentric Castleman Disease (KSHV-MCD) and the KSHV Inflammatory Cytokine Syndrome. Front Microbiol 3:73.

28. Stiller CA. 2007. International patterns of cancer incidence in adolescents. Cancer Treatment Reviews 33:631–645.

29. Chatlynne LG, Ablashi DV. 1999. Seroepidemiology of Kaposi’s sarcoma-associated herpesvirus (KSHV). Seminars in Cancer Biology 9:175–185.

30. Parkin DM, Sitas F, Chirenje M, Stein L, Abratt R, Wabinga H. 2008. Part I: Cancer in Indigenous Africans—burden, distribution, and trends. The Lancet Oncology 9:683–692.

31. Amir H, Kaaya EE, Manji KP, Kwesigabo G, Biberfeld P. 2001. Kaposi’s sarcoma before and during a human immunodeficiency virus epidemic in Tanzanian children: The Pediatric Infectious Disease Journal 20:518–521.

32. Bechtel JT, Liang Y, Hvidding J, Ganem D. 2003. Host Range of Kaposi’s Sarcoma-Associated Herpesvirus in Cultured Cells. JVI 77:6474–6481.

33. Hahn AS, Desrosiers RC. 2013. Rhesus Monkey Rhadinovirus Uses Eph Family Receptors for Entry into B Cells and Endothelial Cells but Not Fibroblasts. PLOS Pathogens 9:e1003360.

34. Dollery SJ. 2019. Towards Understanding KSHV Fusion and Entry. Viruses 11:1073.

35. Hahn AS, Kaufmann JK, Wies E, Naschberger E, Panteleev-Ivlev J, Schmidt K, Holzer A, Schmidt M, Chen J, König S, Ensser A, Myoung J, Brockmeyer NH, Stürzl M, Fleckenstein B, Neipel F. 2012. The ephrin receptor tyrosine kinase A2 is a cellular receptor for Kaposi’s sarcoma–associated herpesvirus. Nat Med 18:961–966.

36. Großkopf AK, Ensser A, Neipel F, Jungnickl D, Schlagowski S, Desrosiers RC, Hahn AS. 2018. A conserved Eph family receptor-binding motif on the gH/gL complex of Kaposi’s sarcoma-associated herpesvirus and rhesus monkey rhadinovirus. 2. PLoS Pathog 14:e1006912.

37. Großkopf AK, Schlagowski S, Hörnich BF, Fricke T, Desrosiers RC, Hahn AS. 2019. EphA7 Functions as Receptor on BJAB Cells for Cell-to-Cell Transmission of the Kaposi’s Sarcoma-Associated Herpesvirus and for Cell-Free Infection by the Related Rhesus Monkey Rhadinovirus. J Virol 93:e00064–19, /jvi/93/15/JVI.00064-19.atom.

38. Großkopf AK, Schlagowski S, Fricke T, Ensser A, Desrosiers RC, Hahn AS. 2021. Plxdc family members are novel receptors for the rhesus monkey rhadinovirus (RRV). PLoS Pathog 17:e1008979.

39. Hahn A, Birkmann A, Wies E, Dorer D, Mahr K, Sturzl M, Titgemeyer F, Neipel F. 2009. Kaposi’s Sarcoma-Associated Herpesvirus gH/gL: Glycoprotein Export and Interaction with Cellular Receptors. J Virol 83:396–407.

40. Akula SM, Pramod NP, Wang F-Z, Chandran B. 2002. Integrin α3β1 (CD 49c/29) Is a Cellular Receptor for Kaposi’s Sarcoma-Associated Herpesvirus (KSHV/HHV-8) Entry into the Target Cells. Cell 108:407–419.

41. Garrigues HJ, DeMaster LK, Rubinchikova YE, Rose TM. 2014. KSHV attachment and entry are dependent on αVβ3 integrin localized to specific cell surface microdomains and do not correlate with the presence of heparan sulfate. Virology 0:118–133.

42. Raghu H, Sharma-Walia N, Veettil MV, Sadagopan S, Chandran B. 2009. Kaposi’s Sarcoma-Associated Herpesvirus Utilizes an Actin Polymerization-Dependent Macropinocytic Pathway To Enter Human Dermal Microvascular Endothelial and Human Umbilical Vein Endothelial Cells. JVI 83:4895–4911.

43. Akula SM, Naranatt PP, Walia N-S, Wang F-Z, Fegley B, Chandran B. 2003. Kaposi’s Sarcoma-Associated Herpesvirus (Human Herpesvirus 8) Infection of Human Fibroblast Cells Occurs through Endocytosis. JVI 77:7978–7990.

44. Zhang W, Zhou F, Greene W, Gao S-J. 2010. Rhesus Rhadinovirus Infection of Rhesus Fibroblasts Occurs through Clathrin-Mediated Endocytosis. Journal of Virology 84:11709–11717.

45. Pertel PE. 2002. Human Herpesvirus 8 Glycoprotein B (gB), gH, and gL Can Mediate Cell Fusion. JVI 76:4390–4400.

46. Chen J, Schaller S, Jardetzky TS, Longnecker R. 2020. Epstein-Barr Virus gH/gL and Kaposi’s Sarcoma-Associated Herpesvirus gH/gL Bind to Different Sites on EphA2 To Trigger Fusion. J Virol 94:e01454–20, /jvi/94/21/JVI.01454-20.atom.

47. Myoung J, Ganem D. 2011. Infection of Lymphoblastoid Cell Lines by Kaposi’s Sarcoma-Associated Herpesvirus: Critical Role of Cell-Associated Virus. Journal of Virology 85:9767–9777.

48. Jarousse N, Chandran B, Coscoy L. 2008. Lack of Heparan Sulfate Expression in B-Cell Lines: Implications for Kaposi’s Sarcoma-Associated Herpesvirus and Murine Gammaherpesvirus 68 Infections. Journal of Virology 82:12591–12597.

49. Dollery SJ, Santiago-Crespo RJ, Kardava L, Moir S, Berger EA. 2013. Efficient infection of a human B cell line with cell-free Kaposi’s sarcoma-associated herpesvirus. J Virol https://doi.org/10.1128/JVI.03063-13.

50. Stoltz M, Klingström J. 2010. Alpha/Beta Interferon (IFN-α/β)-Independent Induction of IFN-λ1 (Interleukin-29) in Response to Hantaan Virus Infection. JVI 84:9140–9148.

51. Tissari J, Sirén J, Meri S, Julkunen I, Matikainen S. 2005. IFN-α Enhances TLR3-Mediated Antiviral Cytokine Expression in Human Endothelial and Epithelial Cells by Up-Regulating TLR3 Expression. J Immunol 174:4289–4294.

52. Thube MM, Shil P, Kasbe R, Patil AA, Pawar SD, Mullick J. 2018. Differences in Type I interferon response in human lung epithelial cells infected by highly pathogenic H5N1 and low pathogenic H11N1 avian influenza viruses. Virus Genes 54:414–423.

53. Spence JS, He R, Hoffmann H-H, Das T, Thinon E, Rice CM, Peng T, Chandran K, Hang HC. 2019. IFITM3 directly engages and shuttles incoming virus particles to lysosomes. 3. Nat Chem Biol 15:259–268.

54. Zhao X, Guo F, Liu F, Cuconati A, Chang J, Block TM, Guo J-T. 2014. Interferon induction of IFITM proteins promotes infection by human coronavirus OC43. 18. Proceedings of the National Academy of Sciences 111:6756–6761.

55. Winkler M, Wrensch F, Bosch P, Knoth M, Schindler M, Gärtner S, Pöhlmann S. 2019. Analysis of IFITM-IFITM Interactions by a Flow Cytometry-Based FRET Assay. 16. IJMS 20:3859.

56. Rahman K, Coomer CA, Majdoul S, Ding SY, Padilla-Parra S, Compton AA. 2020. Homology-guided identification of a conserved motif linking the antiviral functions of IFITM3 to its oligomeric state. eLife 9:e58537.

57. Sanjana NE, Shalem O, Zhang F. 2014. Improved vectors and genome-wide libraries for CRISPR screening. Nat Meth 11:783–784.

58. Tang Y-W, Johnson JE, Browning PJ, Cruz-Gervis RA, Davis A, Graham BS, Brigham KL, Oates JA, Loyd JE, Stecenko AA. 2003. Herpesvirus DNA Is Consistently Detected in Lungs of Patients with Idiopathic Pulmonary Fibrosis. Journal of Clinical Microbiology 41:2633–2640.

59. Lin Y-C, Boone M, Meuris L, Lemmens I, Van Roy N, Soete A, Reumers J, Moisse M, Plaisance S, Drmanac R, Chen J, Speleman F, Lambrechts D, Van de Peer Y, Tavernier J, Callewaert N. 2014. Genome dynamics of the human embryonic kidney 293 lineage in response to cell biology manipulations. 1. Nature Communications 5:4767.

60. Stürzl M, Gaus D, Dirks WG, Ganem D, Jochmann R. 2013. Kaposi’s sarcoma-derived cell line SLK is not of endothelial origin, but is a contaminant from a known renal carcinoma cell line. Int J Cancer 132:1954–1958.

61. Myoung J, Ganem D. 2011. Generation of a doxycycline-inducible KSHV producer cell line of endothelial origin: Maintenance of tight latency with efficient reactivation upon induction. Journal of Virological Methods 174:12–21.

62. Jia R, Xu F, Qian J, Yao Y, Miao C, Zheng Y-M, Liu S-L, Guo F, Geng Y, Qiao W, Liang C. 2014. Identification of an endocytic signal essential for the antiviral action of IFITM3: Endocytosis of IFITM3 and its antiviral activity. Cell Microbiol 16:1080–1093.

63. Chesarino NM, McMichael TM, Hach JC, Yount JS. 2014. Phosphorylation of the Antiviral Protein Interferon-inducible Transmembrane Protein 3 (IFITM3) Dually Regulates Its Endocytosis and Ubiquitination. Journal of Biological Chemistry 289:11986–11992.

64. Zani A, Zhang L, McMichael TM, Kenney AD, Chemudupati M, Kwiek JJ, Liu S-L, Yount JS. 2019. Interferon-induced transmembrane proteins inhibit cell fusion mediated by trophoblast syncytins. Journal of Biological Chemistry 294:19844–19851.

65. Dutta D, Chakraborty S, Bandyopadhyay C, Valiya Veettil M, Ansari MA, Singh VV, Chandran B. 2013. EphrinA2 Regulates Clathrin Mediated KSHV Endocytosis in Fibroblast Cells by Coordinating Integrin-Associated Signaling and c-Cbl Directed Polyubiquitination. PLoS Pathog 9:e1003510.

66. Perreira JM, Chin CR, Feeley EM, Brass AL. 2013. IFITMs Restrict the Replication of Multiple Pathogenic Viruses. Journal of Molecular Biology 425:4937–4955.

67. Hunt CL, Lennemann NJ, Maury W. 2012. Filovirus Entry: A Novelty in the Viral Fusion World. Viruses 4:258–275.

68. Vidak E, Javoršek U, Vizovišek M, Turk B. 2019. Cysteine Cathepsins and their Extracellular Roles: Shaping the Microenvironment. Cells 8:264.

69. Winkler M, Gärtner S, Wrensch F, Krawczak M, Sauermann U, Pöhlmann S. 2017. Rhesus macaque IFITM3 gene polymorphisms and SIV infection. 3. PLoS ONE 12:e0172847.

70. Wilkins J, Zheng Y-M, Yu J, Liang C, Liu S-L. 2016. Nonhuman Primate IFITM Proteins Are Potent Inhibitors of HIV and SIV. PLoS ONE 11:e0156739.

71. Hickford D, Frankenberg S, Shaw G, Renfree MB. 2012. Evolution of vertebrate interferon inducible transmembrane proteins. BMC Genomics 13:155.

72. Zhang Z, Liu J, Li M, Yang H, Zhang C. 2012. Evolutionary Dynamics of the Interferon-Induced Transmembrane Gene Family in Vertebrates. PLoS ONE 7:e49265.

73. Kummer S, Avinoam O, Kräusslich H-G. 2019. IFITM3 Clusters on Virus Containing Endosomes and Lysosomes Early in the Influenza A Infection of Human Airway Epithelial Cells. 6. Viruses 11:548.

74. Ren L, Du S, Xu W, Li T, Wu S, Jin N, Li C. 2020. Current Progress on Host Antiviral Factor IFITMs. Front Immunol 11:543444.

75. Buchrieser J, Dufloo J, Hubert M, Monel B, Planas D, Rajah MM, Planchais C, Porrot F, Guivel-Benhassine F, Van der Werf S, Casartelli N, Mouquet H, Bruel T, Schwartz O. 2020. Syncytia formation by SARS-CoV-2-infected cells. EMBO J 39.

76. Hörnich BF, Großkopf AK, Schlagowski S, Tenbusch M, Kleine-Weber H, Neipel F, Stahl-Hennig C, Hahn AS. 2021. SARS-CoV-2 and SARS-CoV Spike-Mediated Cell-Cell Fusion Differ in Their Requirements for Receptor Expression and Proteolytic Activation. J Virol 95.

77. Ensser A, Yasuda K, Lauer W, Desrosiers RC, Hahn AS. 2020. Rhesus Monkey Rhadinovirus Isolated from Hemangioma Tissue. Microbiol Resour Announc 9.

78. Tischer BK, von Einem J, Kaufer B, Osterrieder N. 2006. Two-step red-mediated recombination for versatile high-efficiency markerless DNA manipulation in Escherichia coli. BioTechniques 40:191–197.

